# Ca_V_2.1 α_1_ subunit motifs that control presynaptic Ca_V_2.1 subtype abundance are distinct from Ca_V_2.1 preference

**DOI:** 10.1101/2023.04.28.538778

**Authors:** Jianing Li, Priyadharishini Veeraraghavan, Samuel M. Young

## Abstract

Presynaptic voltage-gated Ca^2+^ channels (Ca_V_) subtype abundance at mammalian synapses regulates synaptic transmission in health and disease. In the mammalian central nervous system, most presynaptic terminals are Ca_V_2.1 dominant with a developmental reduction in Ca_V_2.2 and Ca_V_2.3 levels, and Ca_V_2 subtype levels are altered in various diseases. However, the molecular mechanisms controlling presynaptic Ca_V_2 subtype levels are largely unsolved. Since the Ca_V_2 α_1_ subunit cytoplasmic regions contain varying levels of sequence conservation, these regions are proposed to control presynaptic Ca_V_2 subtype preference and abundance. To investigate the potential role of these regions, we expressed chimeric Ca_V_2.1 α_1_ subunits containing swapped motifs with the Ca_V_2.2 and Ca_V_2.3 α_1_ subunit on a Ca_V_2.1/Ca_V_2.2 null background at the calyx of Held presynaptic terminal. We found that expression of Ca_V_2.1 α_1_ subunit chimeras containing the Ca_V_2.3 loop II-III region or cytoplasmic C-terminus (CT) resulted in a large reduction of presynaptic Ca^2+^ currents compared to the Ca_V_2.1 α_1_ subunit. However, the Ca^2+^ current sensitivity to the Ca_V_2.1 blocker Agatoxin-IVA, was the same between the chimeras and the Ca_V_2.1 α_1_ subunit. Additionally, we found no reduction in presynaptic Ca^2+^ currents with Ca_V_2.1/2.2 cytoplasmic CT chimeras. We conclude that the motifs in the Ca_V_2.1 loop II-III and CT do not individually regulate Ca_V_2.1 preference, but these motifs control Ca_V_2.1 levels and the Ca_V_2.3 CT contains motifs that negatively regulate presynaptic Ca_V_2.3 levels. We propose that the motifs controlling presynaptic Ca_V_2.1 preference are distinct from those regulating Ca_V_2.1 levels and may act synergistically to impact pathways regulating Ca_V_2.1 preference and abundance.

**Key points summary:**

- Presynaptic Ca_V_2 subtype abundance regulates neuronal circuit properties, however the mechanisms regulating presynaptic Ca_V_2 subtype abundance and preference remains enigmatic.
- The Ca_V_ α_1_ subunit determines subtype and contains multiple motifs implicated in regulating presynaptic subtype abundance and preference.
- The Ca_V_2.1 α_1_ subunit domain II-III loop and cytoplasmic C-terminus are positive regulators of presynaptic Ca_V_2.1 abundance but do not regulate preference.
- The Ca_V_2.3 α_1_ subunit cytoplasmic C-terminus negatively regulates presynaptic Ca_V_2 subtype abundance but not preference while the Ca_V_2.2 α_1_ subunit cytoplasmic C-terminus is not a key regulator of presynaptic Ca_V_2 subtype abundance or preference.
- The Ca_V_2 α_1_ subunit motifs determining the presynaptic Ca_V_2 preference are distinct from abundance.

## Introduction

At mammalian central nervous system (CNS) synapses, Ca_V_2.1, Ca_V_2.2, and Ca_V_2.3, are key determinants of Ca^2+^ entry that controls synaptic transmission and neuronal circuit function (Dolphin & Lee, 2020; Young & Veeraraghavan, 2021). Among the three Ca_V_2 subtypes, presynaptic Ca_V_2.1 levels are dominant compared to Ca_V_2.2 and Ca_V_2.3 (Iwasaki & Takahashi, 1998; Iwasaki *et al*., 2000; Nanou & Catterall, 2018; Dolphin & Lee, 2020). During neuronal circuit maturation, action-potential (AP) evoked synaptic transmission becomes increasingly Ca_V_2.1-dependent due to a selective presynaptic reduction of Ca_V_2.2 and Ca_V_2.3 (Iwasaki & Takahashi, 1998; Eggermann *et al*., 2011; Young & Veeraraghavan, 2021). In addition, presynaptic Ca_V_2 subtype levels vary between synapses and are altered in various diseases (Dolphin & Lee, 2020; Young & Veeraraghavan, 2021). Since Ca_V_2 subtypes differentially control Ca^2+^ influx, elucidating the molecular mechanisms controlling presynaptic Ca_V_2 subtype abundance are critical to understanding brain function in health and disease.

Although presynaptic Ca_V_2 subtype levels are highly variable, they are distinct between the neuronal soma and presynaptic terminal (Doughty *et al*., 1998). In addition, loss of Ca_V_2.1 leads to increases in presynaptic Ca_V_2.2 levels, however total presynaptic Ca^2+^ currents are reduced (Jun *et al*., 1999; Inchauspe *et al*., 2004; Lubbert *et al*., 2017). Therefore, it is proposed that mechanisms that control presynaptic Ca_V_2 levels and preferences are distinct from those controlling somatic Ca_V_2 levels (Weiss & Zamponi, 2017; Dolphin & Lee, 2020). However, whether the mechanisms that determine presynaptic Ca_V_2 subtype preference and abundance are interrelated or separable are unknown.

The pore-forming Ca_V_2 α_1_ subunit determines presynaptic Ca_V_2 preference and contains multiple cytoplasmic protein interaction motifs (Dolphin & Lee, 2020). Due to varying levels of sequence conservation between the three Ca_V_2 α_1_ subunit isoforms cytoplasmic regions, three regions are implicated in regulating presynaptic Ca_V_2 channel levels. They are: 1) the I-II loop; the primary binding site to the Ca_V_β subunit (AID) 2) the II-III loop; the synprint domain; and 3) the C-terminal region, which contains multiple active zone (AZ) protein binding sites and is frequently mutated in Ca_V_2 channelopathies (Dolphin & Lee, 2020; Young & Veeraraghavan, 2021). Since the loop II-III and cytoplasmic CT regions are not highly conserved between the different Ca_V_2 α_1_ subunit isoforms and are subject to alternative splicing (Simms, 2014), they are speculated to contain motifs that regulate the presynaptic Ca_V_2 subtypes levels and preference. However, the identity of the Ca_V_2 α_1_ subunit motifs that regulate presynaptic Ca_V_2 subtype levels and preference is unclear. Studies from cultured superior cervical ganglion neurons demonstrated that the Ca_V_2.1 loop II-III domain regulates presynaptic targeting (Mochida *et al*., 2003). However, it is not defined if the Ca_V_2.1 loop II-III region is a key determinant of preference or if the II-III region from other Ca_V_2 subtypes regulates abundance and preference. Although motifs in the Ca_V_2.1 CT (aa. 2016-2368) that interact with AZ proteins are dispensable for controlling presynaptic Ca_V_2.1 abundance (Lubbert *et al*., 2017), it is not known if other regions in the Ca_V_2.1 CT control Ca_V_2 subtype abundance and preference. Furthermore, whether the Ca_V_2.2 and Ca_V_2.3 CT regulates abundance and preference is unresolved.

To elucidate the roles of the Ca_V_2 α_1_ subunit loop II-III and CT in regulating presynaptic abundance and preference in a native neuronal circuit, we generated Ca_V_2 α_1_ subunit chimeras: 1) <u>Ca_V_2.1/2.3 chimeras</u>, the Ca_V_2.1 α_1_ subunit II-III loop and/or CT region, were swapped with these regions from the Ca_V_2.3 α_1_ subunit. 2) <u>Ca_V_2.1/2.2 chimeras,</u> Ca_V_2.1 α_1_ subunit CT region was swapped with the Ca_V_2.2 α_1_ subunit CT region. We then used helper-dependent Adenoviral (HdAd) technology and delivery techniques to express Ca_V_2.1 α_1_ subunit chimeras on a Ca_V_2.1/Ca_V_2.2 null background at the postnatal days (P) 9–11 calyx of Held using a *Cacna1a*/*Cacna1b flox/flox* (Ca_V_2.1 Ca_V_2.2 CKO) mouse model. The calyx of Held is a large glutamatergic presynaptic terminal in the auditory brainstem that at this developmental stage contains multiple Ca_V_2 subtypes which is similar to the majority of presynaptic terminals (Iwasaki *et al*., 2000; Young & Veeraraghavan, 2021).

Using direct presynaptic Ca^2+^ current recordings at the calyx of Held, we found that ablation of Ca_V_2.1 and Ca_V_2.2 resulted in a significant reduction in total presynaptic Ca^2+^ currents and were insensitive ω-Agatoxin-IVA (Aga), a Ca_V_2.1 specific blocker, and ω-Conotoxin GVIA (Cono), a Ca_V_2.2 specific blocker, and Nifedipine a Ca_V_1 specific blocker, but were partially sensitive to SNX-482, a Ca_V_2.3 specific blocker. Expression of Ca_V_2.1/2.3 chimeras with single or combined region swaps resulted in a significant reduction in peak Ca^2+^ current amplitudes. However, the remaining Ca^2+^ currents with the single swap Ca_V_2.1/2.3 chimeras exhibited a similar proportion of Aga sensitive currents as the full transcript (FT) Ca_V_2.1 α_1_ subunit, while the combined swap had a significant increase in Aga insensitive Ca^2+^ currents. In addition, expression of Ca_V_2.1/2.3 chimeras which replaced the dispensable region of the Ca_V_2.1 CT (aa. 2016-2368) with corresponding Ca_V_2.3 CT region resulted in a reduction in Ca^2+^ currents. Finally, we found that expression of Ca_V_2.1/2.2 CT chimeras revealed no reduction in presynaptic Ca^2+^ currents.

Based on our data, we conclude that individual motifs in the Ca_V_2.1 α_1_ subunit loop II-III or CT are positive regulators of presynaptic Ca_V_2.1 levels but do not individually control Ca_V_2.1 preference. Furthermore, we conclude the Ca_V_2.3 CT contains motifs that negatively regulate presynaptic Ca_V_2.3 levels. Therefore, we propose that the motifs that determine presynaptic Ca_V_2.1 abundance are distinct from those regulating Ca_V_2.1 preference and these motifs may synergistically impact pathways that regulate Ca_V_2.1 preference and levels.

## Methods and materials

### Ethical approval

All procedures were performed according to the animal welfare law of the University of Iowa Office of Animal Resources (OAR) and Institutional Animals Care and Use Committee (IACUC, D16-00009 (A3021-01)). Mice are maintained at The University of Iowa Animal Care Facility which is approved under NIH assurance A3021-01 and certified by the State of Iowa for use of living animals. Animals were housed with 12 h light/dark cycle and *ab libitum* food/water supply. Animals of both sexes at postnatal day 9-11 were used for all experiments. All available measures were taken to minimize animal pain and suffering.

### EXPERIMENTAL DESIGN

#### Generation of *Cacna1a*/*Cacna1b* flox/flox animal

##### Cacna1a fl/fl mice

*Cacna1a^fl/fl^* mice were generated as follows. The Cacna1a^tm1a(EUCOMM)Hmgu^ targeting vector (EUMMCR) was electroporated into the JM8A3 ES-cells derived from C57BL/6N mice and clones were selected and PCR verified. The neo gene was removed by crossing to FLPo mice carrying the FLP recombinase gene under the control of the CAG promoter (C57BL/6N-Tg (CAG-Flpo)1Afst/Mmucd, stock 036512-UCD. (Mouse Biology Program KOMP). Genotyping of *Cacna1a* flox mice by PCR was carried out by the following primers,

**754** 5’ GGAATCCCAAGTGCGTGAGTCTCC 3’

**887** 5’ TCAGCCCCTTCTCTTTAATCTGACCC 3’

*Cacna1a^fl/fl^* mice were backcrossed to C57BL6/J background.

##### Cacna1b^fl/fl^ mice

*Cacna1b^fl/fl^ mice* (*C*57BL/6N-Cacna1b^tm1c(KOMP)Wtsi^/H) were purchased from MRC Harwell Institute. Genotyping of *Cacna1b^fl/fl^*was carried out using the following primers,

**Cacna1b-5arm-WTF** 5’ TTGGTGTCCTGTTCTCCACA 3’

**Cacna1b-Crit-WTR** 5’ CAGGGAATCCTGGGAAGTAA 3’

**5mut-R1** 5’ GAACTTCGGAATAGGAACTTCG 3’

*Cacna1b^fl/fl^ mice* were backcrossed to the C57BL6/J background.

*Cacna1a^fl/fl^, Cacna1b^fl/fl^* mice

*Cacna1a^fl/fl^, Cacna1b^fl/fl^* mice were generated by crossbreeding *Cacna1a^fl/fl^* with *Cacna1b^fl/fl^* to homozygosity. Genotyping was performed by PCR.

### Stereotactic P1 surgery

Stereotactic surgery was performed at P1 as described previously (Radulovic *et al*., 2020). Briefly, *Cacna1a^fl/fl^*, *Cacna1b^fl/fl^*mice were anesthetized by hypothermia, using ice bath for 5min. Subsequently, ∼2 µl HdAd in storage buffer (in Mm: 10 HEPES, 250 sucrose, 1 MgCl_2_ at pH: 7.4 and 6.6% mannitol) was injected at 1 μl/min into the cochlear nucleus (CN) using pulled glass pipettes with a 20 μm opening (Sutter Instrument, P-97, U.S.A.). Two viral vectors, expressing Cre and Ca_V_2 α_1_ hybrid constructs were co-injected. The total viral particle delivered did not exceed 2*10^9^ viral particles. To dissipate pressure after injection, the needle was slowly removed. Animals were then placed under a warming lamp. Upon full recovery, visual inspection and tail pinch responsiveness, animals were returned to their respective cage with the dam. If animals did not recover from the surgery, they were immediately euthanized. In the case of runting or if animals were found in distress at later time points the veterinary staff was consulted and their recommendation for further action was followed.

### DNA constructs and recombinant viral vector production

#### Ca_V_2.1 chimeras

Codon optimized (GENEart) Ca_V_2.1 α_1_ subunit cDNA Cacna1a (*Mus musculus*, Accession No. NP_031604.3), Ca_V_2.2 α_1_ subunit cDNA Cacna1b (*Mus musculus,* Accession No. NP_001035993.1) and Ca_V_2.3 α_1_ subunit cDNA Cacna1e (*Mus musculus*, Accession No. NP_033912.2) were used to generate Ca_V_2 α_1_ hybrid constructs. All Ca_V_2 hybrid constructs were sequenced verified. Amino acid alignment of above constructs were performed under Clustal Omega function using the MegAlign Pro 17 program implemented in the Lasergene software package (v.17, DNASTAR). The following Ca_V_2 hybrid constructs were created.

#### Ca_V_2.1/2.3 II-III loop

Ca_V_2.1 α_1_ subunit aa. 719-1210 replaced with Ca_V_2.3 α_1_ subunit aa. 707-1151

#### Ca_V_2.1/2.3 CT

Ca_V_2.1 α_1_ subunit aa. 1933-2368 replaced with Ca_V_2.3 α_1_ subunit aa. 1897-2273

#### Ca_V_2.1/2.3 II-III loop CT

Ca_V_2.1 α_1_ subunit aa. 719-1210 replaced with Ca_V_2.3 α_1_ subunit aa. 707-1151 and Ca_V_2.1 α_1_ subunit aa. 1933-2368 replaced with Ca_V_2.3 α_1_ subunit aa. 1897-2273

#### Ca_V_2.1/2.2 CT

Ca_V_2.1 α_1_ subunit aa. 1933-2368 replaced by Ca_V_2.2 α_1_ subunit aa. 1870-2327

#### Ca_V_2.1/2.3 CT1

Ca_V_2.1 α_1_ subunit aa. 2016-2368 replaced with Ca_V_2.3 α_1_ subunit 1952-2273

#### Ca_V_2.1/2.3 CT2

Ca_V_2.1 α_1_ subunit aa. 1967-2368 replaced by Ca_V_2.3 α_1_ subunit 1929-2273

For presynaptic Ca^2+^ currents recordings in acute brain slices, these constructs were individually cloned into the EcoRI and NotI sites of the 470 bp human synapsin (hSyn) neurospecific expression cassette (Montesinos *et al*., 2016; Dong *et al*., 2018). For Ca^2+^ currents recordings in HEK293T cells, these constructs were individually cloned into the EcoRI and NotI sites of the 463bp human CMV promoter expression cassette (Tarabichi *et al*., 2023).

### HdAd production

Helper-dependent Adenoviral vector (HdAd) (Cre) which co-expresses Cre recombinase (Cre and EGFP or mCherry under separate neurospecific expression cassettes driven by the 470 bp human synapsin promoter (hSyn) was produced as previously described (Montesinos *et al*., 2016; Dong *et al*., 2018). To produce HdAd Ca_V_2 chimeric viruses we followed standard protocols (Montesinos *et al*., 2016; Lubbert *et al*., 2017). Briefly, Ca_V_2 α_1_ chimera cDNA constructs driven by hSyn promoter expression cassette or a high-level overexpression pUNISHER (Pun) cassette were then cloned into the AscI site, pdelta23E4 which contains a separate neuro-specific expression of mCherry or myristolated EGFP (mEGFP) markers (Montesinos *et al*., 2011). Production of HdAd was carried out as previously described (Palmer & Ng, 2005; Montesinos *et al*., 2016). The HdAd plasmid was linearized with PmeI and then transfected (Profection Mammalian Transfection System, Promega, Madison, WI, USA) into 116 producer cells. Helper virus (HV) was added the following day. Forty-eight hours post infection, cells were subjected to three freeze/thaw cycles and lysate was collected. The HdAd vectors were amplified for total of five serial coinfections from 3×6 cm tissue culture dishes followed by one 15 cm dish and finally 30×15 cm dishes of 116 cells (confluence ∼90%). HdAd was purified by CsCl ultracentrifugation. HdAd was stored at -80ᵒC in storage buffer (10 mM Hepes, 1 mM MgCl_2_, 250 mM Sucrose, pH 7.4).

### HdAd titering and quality control

HdAds were titered by qPCR as described previously (Lubbert *et al*., 2019a). The purified reference virus (RV); Ad5-CMV-EGFP (UNC, Gene Therapy Center, Chapel Hill, North Carolina) was titered by endpoint dilution and by transducing units, as described previously (Bewig & Schmidt, 2000). Human Adenovirus 5 Reference Material (ARM), (ATCC, VR-1516). To titer the HdAd by qPCR, adenoviral gDNA from HdAd, RV and ARM was purified from 6 cm plates. Plates were seeded with 3.75 x 10^6^ HEK293 cells in DMEM with 10% FBS and coinfected with HdAd plus RV on the following day at a multiplicity of 100 vp/cell. The cells were incubated for 1 hr at 37°C with 5% CO_2_ and washed to remove unabsorbed virions. Cells were harvested 4 hr post-infection, then washed with 1 mL PBS and centrifuged for 5 min at 4°C. Pellets were resuspended in 100 mM Tris-HCl, pH 8, lysed by freeze/thaw cycles and treated with 20 U/mL Benzonase for 10 min. Subsequently, lysate was incubated in Proteinase K-SDS solution for 3 hr at 50°C. After protein precipitation with 1.65 M NaCl on ice, DNA was precipitated from supernatants with isopropanol and 3 M sodium acetate, pH 5.2. Samples were mixed well and incubated for 1 hr at -80°C then spun at 4°C for 30 min. Pellets were washed with cold ethanol, dried and resuspended in 10 mM Tris-Cl, pH 8.0. A plasmid standard containing three specific DNA sequences; Ad5 E1a (replication-competent Adenovirus (RCA) detection), Ad5 E2B Iva2 (viral DNA polymerase and terminal protein precursor, HV-exclusive), and Ad genomic stuffer sequence (Cosmid 346 and HPRT1 gene, in all HdAd-exclusive) was used. Three specific primer/probe sets were used for these 3 regions in the plasmid for the detection in qPCR:

1. **Ad5 E1a (RCA):** probe: NED-5’-AGCACCCCGGGCACGGTTG-3’-MGBNFQ; fwd: 5’-GGGTGAGGAGTTTGTGTTAGATTATG-3’; rev: 5’-TCCTCCGGTGATAATGACAAGA-3’
2. **E2B Iva2 (HV):** probe: FAM-5’-TGTCTTTCAGTAGCAAGCT-3’-TAMRA, HV fwd primer: 5’-TGGGCGTGGTGCCTAAAA-3’, rev: 5’-GCCTGCCCCTGGCAAT-3’
3. **Genomic stuffer DNA (HdAd):** probe: VIC-5’-AGCCTCTCTCATCTCACAGT -3’-MGBNFQ, fwd: 5’-CCCCGCTACCCCAATCC-3’, rev: 5’-TTAGCTTTTTTGGGTGATTTTTCC-3’.

Sample DNA was diluted 1:10³, 1:10^4^ or 10^5^ in 10 mM Tris-Cl, pH 8.0. For HdAd and HV probes, a 6-point standard curve was constructed with the plasmid standard using 10^7^-10^2^ molecules/well. For RCA detection, two 7-point curves were constructed with 10^7^ - 10 molecules/well (one plasmid standard and one using ARM gDNA spiked with 10^7^ copies of HdAd gDNA). Real-Time PCR parameters were 50°C for 2 min, 95°C for 10 min; 40 cycles of: 95°C for 15 sec, 60°C for 1 min. Outliers and results out of range of standard curve were discarded. Results within range of the standard curve were corrected for their dilution and determined as HdAd vector genomes/mL. All samples were negative for RCA.

To determine the IU copies/mL from purified coinfection samples, averaged results were divided by 3 µL and multiplied by 25 µL giving IU copies/coinfection. Next, this number was divided by the volume in µL of HdAd added to the coinfection giving IU copies/µL then multiplied by 1000 to obtain the i,nfectious HdAd genomes/mL.

Calculation to determine the titer of purified HdAd virus:

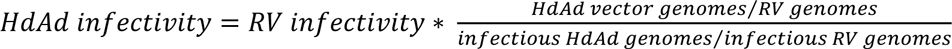

where Ad5-CMV-EGFP is RV, RV infectivity is VP/IU. IU is the average titer of end point dilution and transducing unit assays, HdAd vector genomes and RV genomes are the number of genomes from purified virus as determined by qPCR, and infectious HdAd genomes and infectious RV genomes are the number of genomes in DNA purified from coinfections. The titers of HdAds carry different cDNA constructs are listed as follow, viral particle/ml: Cre EGFP, 1.75*10^12^, Cre mCherry, 2.41*10^12^, syn Ca_V_2.1 FT, 2.40*10^12^, Pun Ca_V_2.1 FT, 2.19*10^12^, Ca_V_2.1/2.3 II-III loop CT, 2.00*10^12^, Ca_V_2.1/2.3 II-III loop, 2.07*10^12^, Syn Ca_V_2.1/2.3 CT, 1.90*10^12^, Pun Ca_V_2.1/2.3 CT, 3.11*10^12^, Ca_V_2.1 delta 2016, 1.40*10^12^, Ca_V_2.1/2.3 CT 2016-2368, 4.70*10^12^, Ca_V_2.1/2.3 CT 1967-2368, 3.29*10^12^, syn Ca_V_2.1/2.2 CT, 3.17*10^12^, Pun Ca_V_2.1/2.2 CT, 9.52*10^12^. Particle to infectious unit ratio was 10:1∼ 50:1. Either syn Cre syn EGFP or syn Cre syn mCherry was used to make sure that the viruses expressing Cre carry different fluorescent protein than the viruses expressing Ca_V_2 constructs.

### Western Blot

To determine expression of our Ca_V_2.1 chimeric constructs, we performed Western Blot analysis on HEK293 cells transduced with our HdAd vectors expressing the Ca_V_2 α_1_ chimeric constructs. To do so, HEK293 cells were infected with our HdAd vectors, and 48 hrs later cells were collected and lysed in RIPA buffer containing: 10 mM Tris-Cl pH8.0, 1mM EDTA, 1% Triton-X, 0.1% sodium deoxycholate, 0.1% SDS, 140 mM NaCl, protease inhibitor (11836153001, Roche). 20 μg of protein lysate were loaded and analyzed by SDS-PAGE followed by immunoblotting: Ca_V_2.1 α_1_ CT antibody (Synaptic Systems, #152 203, rabbit polyclonal) diluted 1:1000 (final concentration 2 μg/ml) and Ca_V_2.1 α_1_ II-III loop antibody (Alomone Labs, ACC-001, rabbit polyclonal) diluted 1:500 (final concentration 1.7 μg/ml) were incubated overnight at 4ᵒC to blot the expression of Ca_V_2.1/2.3 II-III loop hybrid construct, Ca_V_2.3 α_1_ CT antibody (Synaptic Systems, #152 411, mouse monoclonal) diluted 1:500 (final concentration 2 μg/ml) was incubated overnight at 4ᵒC to blot the expression of Ca_V_2.1/2.3 II-III loop CT and Ca_V_2.1/2.3 CT chimeras, Ca_V_2.2 CT antibody (Synaptic Systems, #152 311, mouse monoclonal) diluted 1:1000 (final concentration 2 μg/ml) was incubated overnight at 4ᵒC to detect the expression of Ca_V_2.1/2.2 CT chimera. β-actin antibody (sigma, A2228, mouse monoclonal) diluted 1:5000 (final concentration 0.4 μg /ml) was incubated overnight at 4ᵒC used to blot the expression of β-actin. Goat anti mouse IgG (H+L) poly-HRP (Thermo Fisher, #32230) diluted 1:2000 (final concentration 0.25 μg /ml) and goat anti rabbit IgG (H+L) poly-HRP (Thermo Fisher, #32260) secondary antibody diluted 1:2000 (final concentration 0.25 ng/ml) was incubated at room temperature for 1 hour. Protein bands were detected by chemiluminescence using ECL Western blot detection reagents (Bio-Rad, #170-5060).

### Preparation of acute slices

Acute brain slices were prepared as previously described (Chen *et al*., 2013). Briefly, after decapitation of P9-11 mice of either sex, the brains were immersed in ice-cold low Ca^2+^ artificial cerebrospinal fluid (aCSF) containing (in mM): 125 NaCl, 2.5 KCl, 3 MgCl_2_, 0.1 CaCl_2_, 10 glucose, 25 NaHCO_3_, 1.25 Na_2_HPO_4_, 0.4 L-ascorbic acid, 3 myo-inositol, and 2 Na-pyruvate, pH 7.3-7.4 (310 mosmol/l). aCSF solution was continuously bubbled with 95% O_2_ and 5% CO_2_. Acute brain slices (200 μm thickness) containing MNTB were obtained using a vibrating tissue slicer (Campden 7000 smz-2, Campden Instruments LTD, Loughborough, England). Slices were immediately transferred to standard aCSF (37ᵒC, continuously bubbled with 95% O_2_-5% CO_2_) containing the same chemicals as the slicing buffer except that it contained 1mM MgCl_2_ and 1 mM CaCl_2_. After 1 hour incubation for recovery, slices were transferred to a recording chamber with the same aCSF at room temperature (RT: 25ᵒC).

### Acute slice electrophysiology

In all experiments, slices were continuously perfused with aCSF and visualized by upright microscope (BX51W, Olympus) through 4X air or 60X water-immersion objective (LUMPlanFL N, Olympus, Tokyo, Japan) and a CCD camera (QI-Click, QImaging, Surrey, BC, Canada). Patch clamp recordings were performed by using an EPC 10/2 patch-clamp amplifier (HEKA, Lambrecht, Germany), operated by PatchMaster version 2×90.5 (Harvard Instruments, Holliston, MA, USA). Data were low-pass filtered at 6 kHz and sampled with a rate of 50 kHz. Calyces transduced with HdAd Ca_V_2.1 α_1_ chimeric constructs were identified visually with two co-expressed mEGFP and mCherry markers. To visualize mEGFP and mCherry with light of 470 nm or 560 nm, respectively, a Polychrome V xenon bulb monochromator or a lumen Metal arc lamp was used (TILL Photonics, Grafelfing, Germany).

### Presynaptic Ca^2+^ current recordings

To isolate presynaptic Ca^2+^ currents, aCSF was supplemented with 1 or 1.5 μM tetrodotoxin (TTX, Alomone labs), 100 μM 4-aminopyridin (4-AP, Tocris) and 20 mM tetraethylammonium chloride (TEA, Sigma Aldrich) to block Na^+^ and K^+^ channels. Calyces were whole-cell voltage clamped at -80 mV. Current-voltage (IV) relationships were recorded in the presence of 1 mM CaCl_2_ and 1 mM MgCl_2_ with 1.5 μM TTX while pharmacological isolation of Ca_V_2 subtype components was performed in 2 mM CaCl_2_ and 1 mM MgCl_2_ with 1 μM TTX. To characterize the in the presynaptic terminal, we used 200 nM ω-agatoxin IVA (Alomone labs) for selective inhibition of Ca_V_2.1 channels, 2 μM ω-conotoxin GIVA (Alomone labs) for selective inhibition of Ca_V_2.2 channels, 200 nM SNX-482 (Alomone labs) for selective inhibition of Ca_V_2.3 channels, 10 μM Nifedipine (Alomone labs) for selective inhibition of Ca_V_1 chnannels and supplemented 0.1 mg/mL cytochrome C to prevent toxin absorption. 50μM CdCl_2_ was used for non-selective inhibition of all VGCCs. Presynaptic patch pipette with open tip diameters 4-6 MΩ resistance were pulled from 1.5 mm thin-walled borosilicate glass (Harvard apparatus) and were filled with following internal solution (in mM): 145 Cs-gluconate, 20 TEA-Cl, 10 HEPES, 2 Na_2_-phosphocreatine, 4 MgATP, 0.3 NaGTP, and 0.075 EGTA, pH 7.2, 325-340 mOsm. Pipettes were coated with dental wax to minimize stray capacitance and improve voltage clamp quality. Presynaptic series resistance was less than 15 MΩ and was compensated online to 3 MΩ or less for IV recordings or compensated to 6 MΩ for Ca_V_2 subtype characterization experiments with a time lag of 10 µs by the HEKA amplifier. Leak and capacitive currents were subtracted online with a P/5 routine. Cells with series resistance >15 MΩ and leak currents > 100 pA were excluded from the analysis.

### HEK 293T Cell Culture

Low-passage-number Human embryonic kidney cells transformed with SV40 T-antigen (HEK293T) cells were cultured in DMEM (Thermo Fisher Scientific) with 10% fetal bovine serum at 37°C in a humidified atmosphere with 5% CO_2_. Cells were grown to 70–80% confluence and transiently transfected with 5μg CMV Ca_V_2.1/2.3 chimera constructs, 10μg CMV Ca_V_β4a (Takahashi *et al*., 2004) and 0.25μg CMV mClover to serve as reporter of positive transfection, following manufacturer’s protocols of PolyJet reagent (SignaGen Laboratories).

### HEK293T cells recordings

24-48hrs post transfection, cells were seeded in multiple 35mm dishes. 5-12 hrs post seeding, HEK293T cells were subjected to whole-cell patch clamp recordings (Gandini *et al*., 2014b) The external solution contained the following (mM): 150 Tris, 1 MgCl_2_, and 10 CaCl_2_(Wang *et al*., 2017a). The internal solution was the same as presynaptic Ca^2+^ recording internal solution, but with 5mM EGTA. Patch pipette resistances were ∼3 MΩ and series resistance were compensated 80%∼90% to ∼1MΩ(Gandini *et al*., 2014a). The holding potential was -80mV, and cells were depolarized to reverse potential 46-52mV with 2mV steps to identify the reversal potential, 10ms duration to ensure maximal activation.

### Analysis of electrophysiological data

All data was analyzed by NeuroMatic (Rothman & Silver, 2018) in Igor Pro (version 8.0, Wavemetrics, Portland, OR, USA). Voltage dependence of channel activation was described by both peak and tail currents as functions of voltage. Peak currents were fitted according to a Hodgkin-Huxley formalism with four independent gates assuming a Goldman-Hodgkin-Katz (GHK) open-channel conductance Ƭ: with E_rev_ as reversal potential, V_m_ as half-maximal activation voltage per gate, and k_m_ as the voltage dependence of activation. Tail currents were measured as peaks minus baseline and fitted with a Boltzmann function:

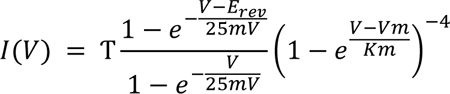

where V_1/2_ represents the half-maximal voltage and k the corresponding slope factor.

For calculation of Q_max_, maximal gating charge of our chimeric constructs in HEK293T cells, we measured as the time integral of the gating currents at reversal potential (Wei *et al*., 1994; Takahashi *et al*., 2004; Wang *et al*., 2017b). The relative *Po* was determined by linear regression, we calculated the relative *Po* by plotting the peak tail current vs. Q_max._ (Wei *et al*., 1994; Takahashi *et al*., 2004; Wang *et al*., 2017b).

### Statistical analysis

All statistical tests were conducted in Prism 6 (GraphPad Software). Sample sizes for all experiments were chosen based on assuming a population with a normal distribution sample size of seven is sufficient to invoke the Central Limit Theorem. All data were tested for normal distribution by performing a Shapiro-Wilk test for normality and variances of all data were estimated and compared using Bartlett’s test. Electrophysiological data were compared with one-way analysis of variance (ANOVA) with a post hoc Dunnett’s test, Ca_V_2.1 FT dataset was used as control group. In Figures and Table, data are reported as mean ± SD. Statistical significance was accepted at *p<0.05; **p<0.01; ***p<0.001; ****p<0.0001.

## Results

### Ca_V_2.1 and Ca_V_2.2 channels can be ablated at the calyx of Held in Ca_V_2.1/Ca_V_2.2 CKO mouse model

Since ablation of Ca_V_2.1 channels results in partial compensation of Ca_V_2 current levels by Ca_V_2.2 channels at the calyx of Held (Inchauspe *et al*., 2004; Lubbert *et al*., 2017), we created a Ca_V_2.1/ Ca_V_2.2 CKO mouse model (Fig. 1A) to allow for spatial and temporal ablation of Ca_V_2.1 and Ca_V_2.2 channels. To determine how presynaptic loss of Ca_V_2.1 and Ca_V_2.2 channels impacted presynaptic Ca_V_2 currents at the calyx of Held, we injected HdAd Cre at P1 into the cochlear nucleus (CN) of Ca_V_2.1/ Ca_V_2.2 CKO mice to create a Ca_V_2.1^-/-^/ Ca_V_2.2^-/-^ calyx of Held. Subsequently we performed whole-cell patch clamp recordings at the P9-11 calyx of Held and characterized the total Ca^2+^ current and current sensitivity to Ca_V_2 subtype-specific blockers: ω-Agatoxin IVA (Aga), Ca_V_2.1-specific, ω-Conotoxin GIVA (Cono), Ca_V_2.2-specific, and Cadmium (Cd^2+^) a non-specific blocker of Ca_V_ currents (Fig. 1D-F, Table 1). Since sequential application of blockers takes ∼10-15 minutes for complete characterization of Ca_V_2 currents, we measured presynaptic Ca^2+^ current rundown at wild-type calyces in the absence of blockers after ∼10-15 minutes (Fig. 1B, C). We found a ∼20-70 pA (2-5%) reduction of presynaptic Ca^2+^ currents (Fig. 1J, Table 1, n=3). Subsequently, analysis of Ca^2+^ currents at the Ca_V_2.1^-/-^/ Ca_V_2.2^-/-^ calyx revealed a significant reduction (wt vs. CKO (pA): -1078 ± 260.9 (n=10) vs. -425.5 ± 219.5 (n=11), p<0.0001) (Fig. 1E, Table 2). We found an almost complete loss of Aga sensitive currents, 809 ± 255.6 (n=4) vs 99.22 ± 53.64 (n=5), p=0.0005; and Cono sensitive currents, 92 ± 142.2 (n=4) vs 50.8 ± 43.1 (n=5), p=0.5532). (Fig1. D and F, Table 1). Therefore, these results demonstrate successful ablation of Ca_V_2.1 and Ca_V_2.2 channels.

**Figure 1.**
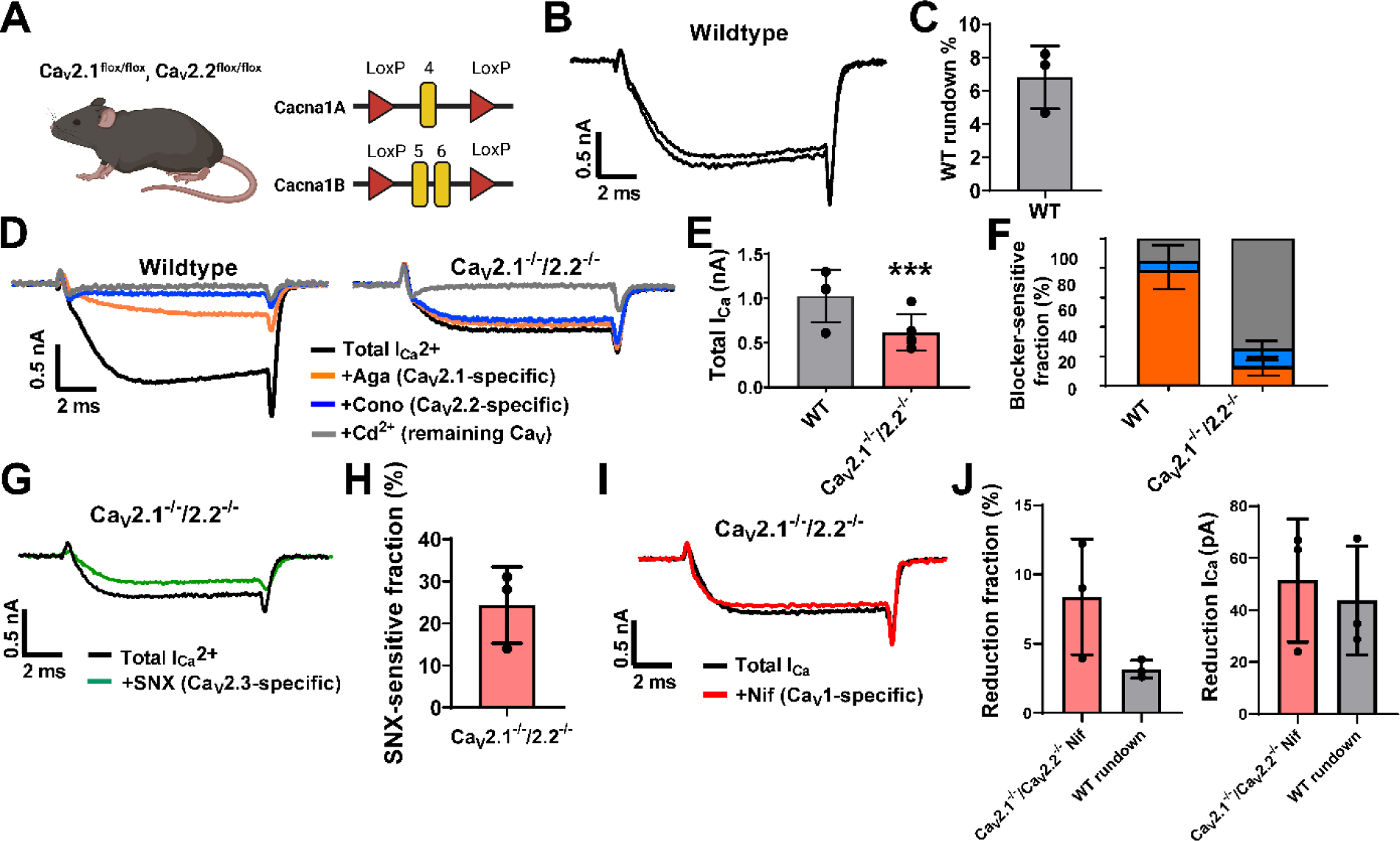
Presynaptic Ca_V_2.1 and Ca_V_2.2 channels can be ablated and the remaining Ca^2+^ currents are compensated by Ca_V_2.3 channels at the calyx of Held. (A) Illustration of CKO mouse model. Exon 4 of *Cacna1a*, and exons 5 and 6 of *Cacna1b* are flanked by LoxP sites. (B-C) Example traces of Ca^2+^ currents and bar graph show rundown fraction of electrophysiological recordings at duration of 10∼15min at wildtype calyx of Held (n=3). (D) Pharmacological isolation of presynaptic Ca_V_2 isoforms in wildtype and Ca_V_2.1^-/-^/ Ca_V_2.2^-/-^ calyces. Ca^2+^ current traces in absence of any blockers (black), after blocking Ca_V_2.1 fraction with 200 nM Aga (orange), after blocking Ca_V_2.2 fraction with 2 mM Cono (blue) and after blocking all Ca_V_ channels with 50 mM Cd^2+^ (gray). (E) Bar graph shows total Ca^2+^ currents in wildtype and Ca_V_2.1^-/-^/ Ca_V_2.2^-/-^ calyces. (F) Relative Ca_V_2 subtype fractions in wildtype, Ca_V_2.1^-/-^/ Ca_V_2.2^-/-^ calyces (n=3-5 for each condition). (G) Ca^2+^ current traces in absence of any blockers (black), after blocking Ca_V_2.3 fraction with 200 nM SNX-482 (green). (H) Bar graph shows SNX-482 sensitive fraction at Ca_V_2.1^-/-^/ Ca_V_2.2^-/-^ calyces (n=3). (I) Ca^2+^ current traces in absence of any blockers (black), after blocking Ca_V_1 fraction with 10 μM Nifedipine (red). (J) Bar graph shows the reduction fraction and Ca^2+^ current amplitudes in Ca_V_2.1^-/-^/ Ca_V_2.2^-/-^ calyces after Nifedipine treatment compared to wildtype rundown fraction.

**Table1.** Cacna1a, *Cacna1b* ^flox/flox^ animal characterization.

| Parameter | Mean | SD | n | P | statistics |
| --- | --- | --- | --- | --- | --- |
| Wildtype animal |  |  |  |  |  |
| Agatoxin sensitive fraction (%) | 79.09 | 14.98 | 4 | control |  |
| Agatoxin sensitive currents (pA) | 809.3 | 255.6 | 4 |  |  |
| Conotoxin sensitive fraction (%) | 7.828 | 10.5 | 4 |  |  |
| Conotoxin sensitive currents (pA) | 92.00 | 142.2 | 4 |  |  |
| Cav2.1<sup>-/-</sup>/ Cav2.2<sup>-/-</sup> |  |  |  |  |  |
| Agatoxin sensitive fraction (%) | 13.64 | 6.24 | 5 | p<0.0001 (****) | Unpaired t test |
| Agatoxin sensitive currents (pA) | 99.22 | 53.64 | 5 | p=0.0005 (***) |  |
| Conotoxin sensitive fraction (%) | 11.92 | 4.722 | 5 | p=0.4571 |  |
| Conotoxin sensitive currents (pA) | 50.80 | 43.10 | 5 | p=0.5532 |  |
| SNX-482 sensitive fraction (%) | 24.33 | 9.074 | 3 |  |  |
| SNX-482 sensitive currents (pA) | 86.65 | 31.26 | 3 |  |  |
| Rundown fraction |  |  |  |  |  |
| Wildtype animal (pA) | 43.70 | 20.85 | 3 | control | Unpaired t test |
| Cav2.1<sup>-/-</sup>/ Cav2.2<sup>-/-</sup> (pA) | 51.40 | 23.78 | 3 |  |  |
| Wildtype animal (%) | 3.167 | 0.6525 | 3 | control | Unpaired t test |
| Cav2.1<sup>-/-</sup>/ Cav2.2<sup>-/-</sup> (%) | 8.393 | 4.169 | 3 | P=0.0985 |  |

**Table2.**
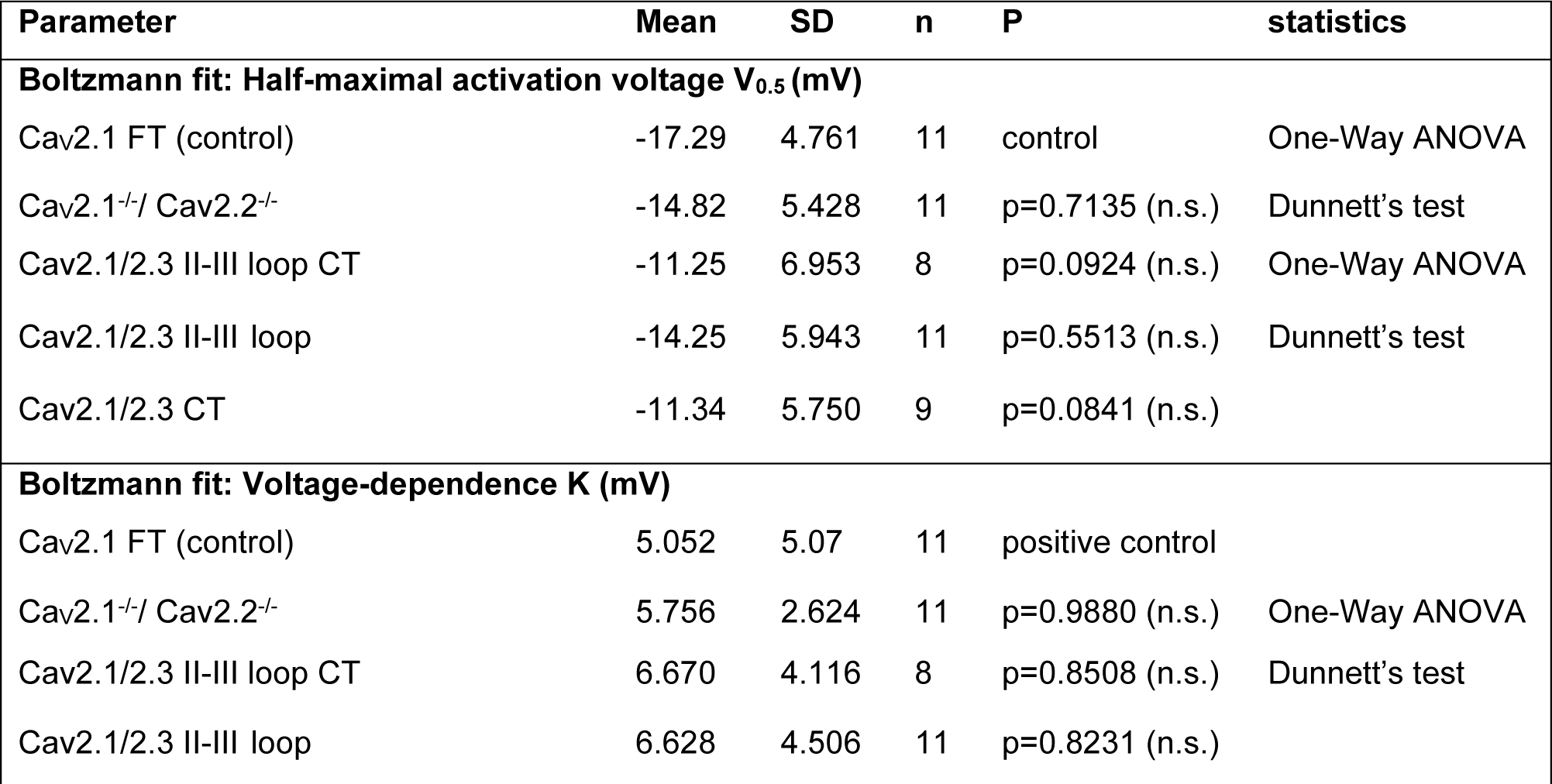
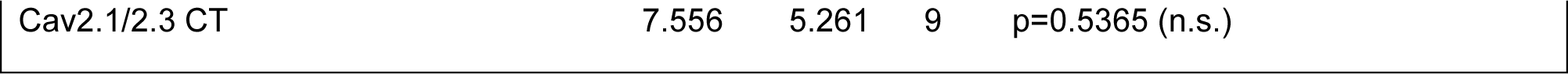
Electrophysiological parameters of IV relations of Ca^2+^ currents.

| Parameter | Mean | SD | n | P | statistics |
| --- | --- | --- | --- | --- | --- |
| **Boltzmann fit: Half-maximal activation voltage V<sub>0.5</sub> (mV)** |  |  |  |  |  |
| Cav2.1 FT (control) | -17.29 | 4.761 | 11 | control | One-Way ANOVA |
| Cav2.1<sup>-/-</sup>/ Cav2.2<sup>-/-</sup> | -14.82 | 5.428 | 11 | p=0.7135 (n.s.) | Dunnett's test |
| Cav2.1/2.3 II-III loop CT | -11.25 | 6.953 | 8 | p=0.0924 (n.s.) | One-Way ANOVA |
| Cav2.1/2.3 II-III loop | -14.25 | 5.943 | 11 | p=0.5513 (n.s.) | Dunnett's test |
| Cav2.1/2.3 CT | -11.34 | 5.750 | 9 | p=0.0841 (n.s.) |  |
| **Boltzmann fit: Voltage-dependence K (mV)** |  |  |  |  |  |
| Cav2.1 FT (control) | 5.052 | 5.07 | 11 | positive control |  |
| Cav2.1<sup>-/-</sup>/ Cav2.2<sup>-/-</sup> | 5.756 | 2.624 | 11 | p=0.9880 (n.s.) | One-Way ANOVA |
| Cav2.1/2.3 II-III loop CT | 6.670 | 4.116 | 8 | p=0.8508 (n.s.) | Dunnett's test |
| Cav2.1/2.3 II-III loop | 6.628 | 4.506 | 11 | p=0.8231 (n.s.) |  |

|  |  |  |  |  |
| --- | --- | --- | --- | --- |
| Cav2.1/2.3 CT | 7.556 | 5.261 | 9 | p=0.5365 (n.s.) |

In the absence of Ca_V_2.1 and Ca_V_2.2, a significant Aga and Cono insensitive presynaptic Ca^2+^ current was present. Since Ca_V_2.3 subtypes can compensate for the loss of Ca_V_2.1 channels (Kaja *et al*., 2006) and Ca_V_1 channels can be found in some presynaptic terminals (Giugovaz-Tropper *et al*., 2011; Young & Veeraraghavan, 2021), it was important to determine if the combined loss of Ca_V_2.1 and Ca_V_2.2 resulted in compensation by Ca_V_2.3 or Ca_V_1 channels. Therefore, we characterized the sensitivity of presynaptic Ca^2+^ current to Ca_V_2.3 and Ca_V_1 specific blockers in the Ca_V_2.1^-/-^/ Ca_V_2.2^-/-^ calyx of Held. First, we used SNX-482, a Ca_V_2.3 selective blocker and found a range of ∼10% to 35% block of presynaptic Ca^2+^ currents (SNX-482 sensitive fraction 24.33 ± 9.074 %) (n=3) (Fig. 1G, H, Table 1). Since there were variable levels of Ca^2+^ current reduction with SNX-482, this indicated the remaining Ca^2+^ currents in the absence of Ca_V_2.1 and Ca_V_2.2 could be mediated by a combination of Ca_V_2.3 variants that are SNX-482 sensitive and SNX-482 insensitive and/or presynaptic Ca_V_1 channels. To determine if Ca_V_1 channels were present, we used nifedipine, a Ca_V_1 selective blocker (Fig. 1I, J) and found presynaptic Ca^2+^ currents were nifedipine insensitive as there was no difference compared to the reduced fraction of Ca^2+^ current due to rundown (Fig. 1I, J, Table 1). Therefore, we conclude that in the absence of Ca_V_2.1 and Ca_V_2.2 channels the compensatory presynaptic Ca^2+^ currents are mediated by mix of SNX-482 sensitive and insensitive Ca_V_2.3 channels.

### The Ca_V_2.1 α_1_ subunit loop II-III domain, and CT are positive regulators of presynaptic Ca_V_2.1 abundance

The amino acid conservation of Ca_V_2.1 α_1_ subunit and Ca_V_2.3 α_1_ subunit loop II-III region is 33.2% and CT region from aa. 1933 onwards is 39.8%, while the loop I-II is 87.3% conserved (Figure 2A, MegAlign Pro 17, Clustal Omega). In fact, the Ca_V_2.3 α_1_ subunit loop II-III lacks a syntaxin I binding site in the synprint region although is modulated by syntaxin IA (Wiser *et al*., 2002) but see (Pereverzev *et al*., 2002), while the Ca_V_2.3 α_1_ subunit CT lacks many of proposed AZ protein binding motifs found in the Ca_V_2.1 CT (Dolphin & Lee, 2020). Since the loop II-III and CT regions are the most divergent, we created Ca_V_2.1/Ca_V_2.3 α_1_ subunit chimeras that swapped the loop II-III and CT domains to determine if these domains regulated presynaptic Ca_V_2.1 abundance and preference (Fig. 2B). We generated three HdAd Ca_V_2.1/2.3 α_1_ subunit chimeric constructs (Figure 2C). There were <u>Ca_V_2.1/2.3 II-III loop</u>, which contains the Ca_V_2.3 α_1_ subunit loop II-III domain (aa. 707 to aa. 1151), <u>Ca_V_2.1/2.3 CT</u>, which contains the cytoplasmic Ca_V_2.3 CT (aa. 1897 to aa. 2273), and <u>Ca_V_2.1/2.3 II-III+CT</u> which contains both the Ca_V_2.3 α_1_ subunit loop II-III and CT (Fig. 3C). To validate that the chimeras were expressed as full-length proteins, we performed Western Blot analysis of HEK293 cells infected with our HdAd Ca_V_2.1/2.3 chimeric constructs. Based on our results, we found that all constructs were successfully expressed (Figure 2D-F). Rescue experiments with an HdAd vector that expressed the Ca_V_2.1 α_1_ subunit FT (Mus musculus NP_031604.3) (HdAd Ca_V_2.1 FT) in the Cav2.1^-/-^ calyx of Held in a *Cacna1a* CKO mouse model has been validated to restore Ca_V_2.1 current amplitudes, subtype levels and synaptic vesicle release rates to similar levels as wild-type calyces (Lubbert *et al*., 2017). To validate our experimental approach, we used expressed the Ca_V_2.1 α_1_ subunit FT using HdAd Ca_V_2.1 FT on the Ca_V_2.1, Ca_V_2.2 CKO mouse line. Our rescue experiments show that HdAd expression of the Ca_V_2.1 FT in the Ca_V_2.1^-/-^/ Ca_V_2.2^-/-^ calyx of Held led to similar maximum Ca^2+^ current amplitudes and voltage-dependent activation as the wild-type calyx of Held. These results validated our experimental approach and confirming previous findings (Figure. 3, Table 2).

**Figure 2.**
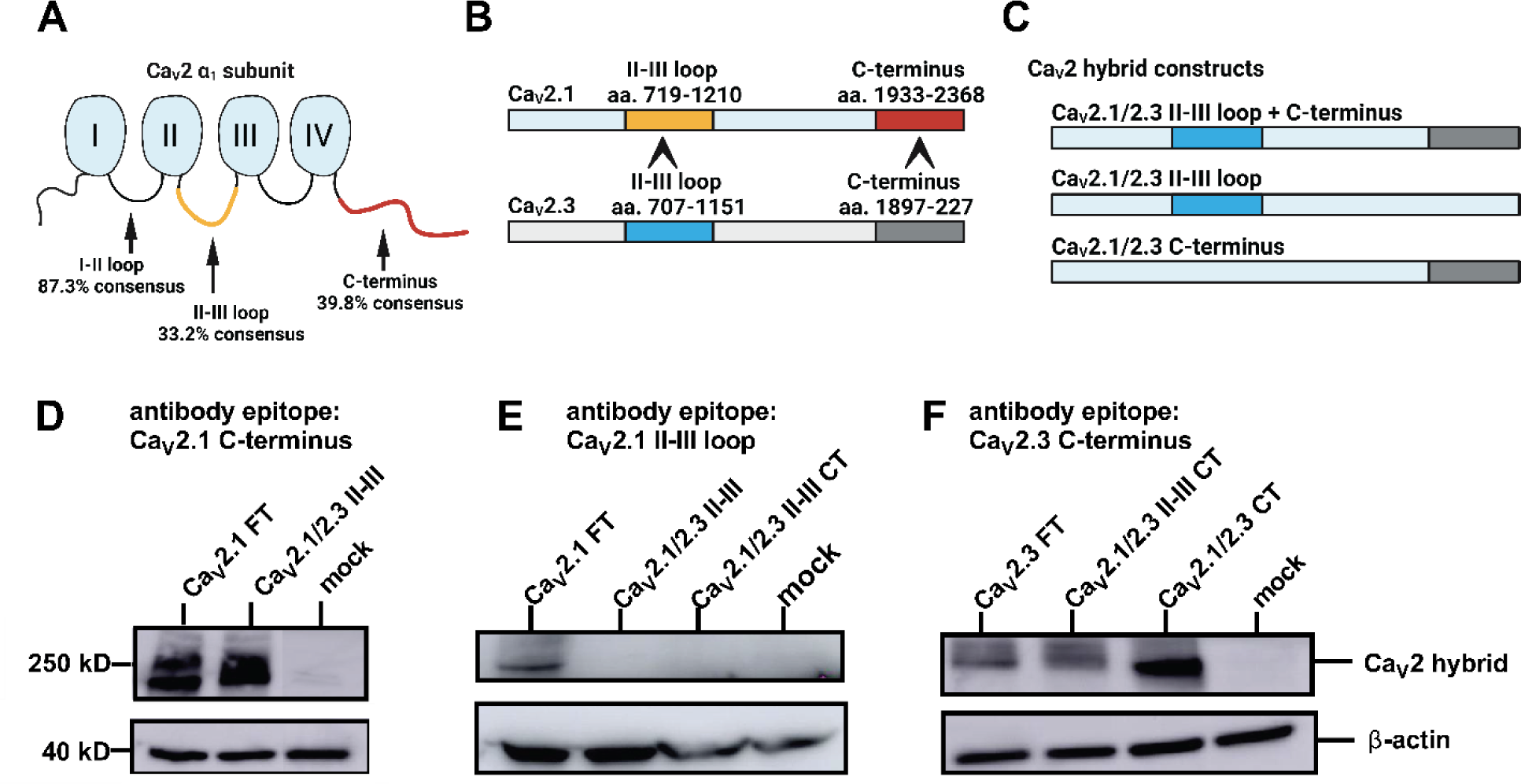
Ca_V_2.1/Ca_V_2.3 Chimeras can be efficiently expressed. Western Blot validated protein expression of Ca_V_2 hybrid constructs. HEK293 cells are infected by HdAd Ca_V_2.1/2.3 chimeric constructs. (A) Illustration of Ca_V_2 α_1_ structure. Four transmembrane domains are highly conserved while the cytoplasmic loops are highly variant between Ca_V_2.1 and Ca_V_2.3, the I-II loop, II-III loop and CT share 87.3%, 33.2% and 39.8% consensus, respectively. (B) Illustration of swapped region between Ca_V_2.1 and Ca_V_2.3 channels, the amino acid numbers are labelled. (C) Illustration of three Ca_V_2.1/2.3 chimeric constructs. Ca_V_2.1 II-III loop and CT are both replaced by Ca_V_2.3 II-III loop and CT in Ca_V_2.1/2.3 II-III loop + CT chimeric construct. Ca_V_2.1 II-III loop is replaced by Ca_V_2.3 II-III loop in Ca_V_2.1/2.3 II-III loop chimeric construct. Ca_V_2.1 CT is replaced by Ca_V_2.3 CT in Ca_V_2.1/2.3 CT chimeric construct. (D) Antibody targeting Ca_V_2.1 CT shows the protein expression of Ca_V_2.1 FT and Ca_V_2.1/2.3 II-III loop. (E) Antibody targeting Ca_V_2.1 II-III loop fails to detect protein expression of Ca_V_2.1/2.3 II-III loop and Ca_V_2.1/2.3 II-III loop CT, indicating that the II-III loop of Ca_V_2.1 is swapped by Ca_V_2.3 II-III loop. (F) Antibody targeting Ca_V_2.3 CT shows the protein expression of Ca_V_2.1/2.3 II-III loop CT and Ca_V_2.1/2.3 CT. Mock, negative control, protein lysis from uninfected HEK293 cells.

**Figure 3.**
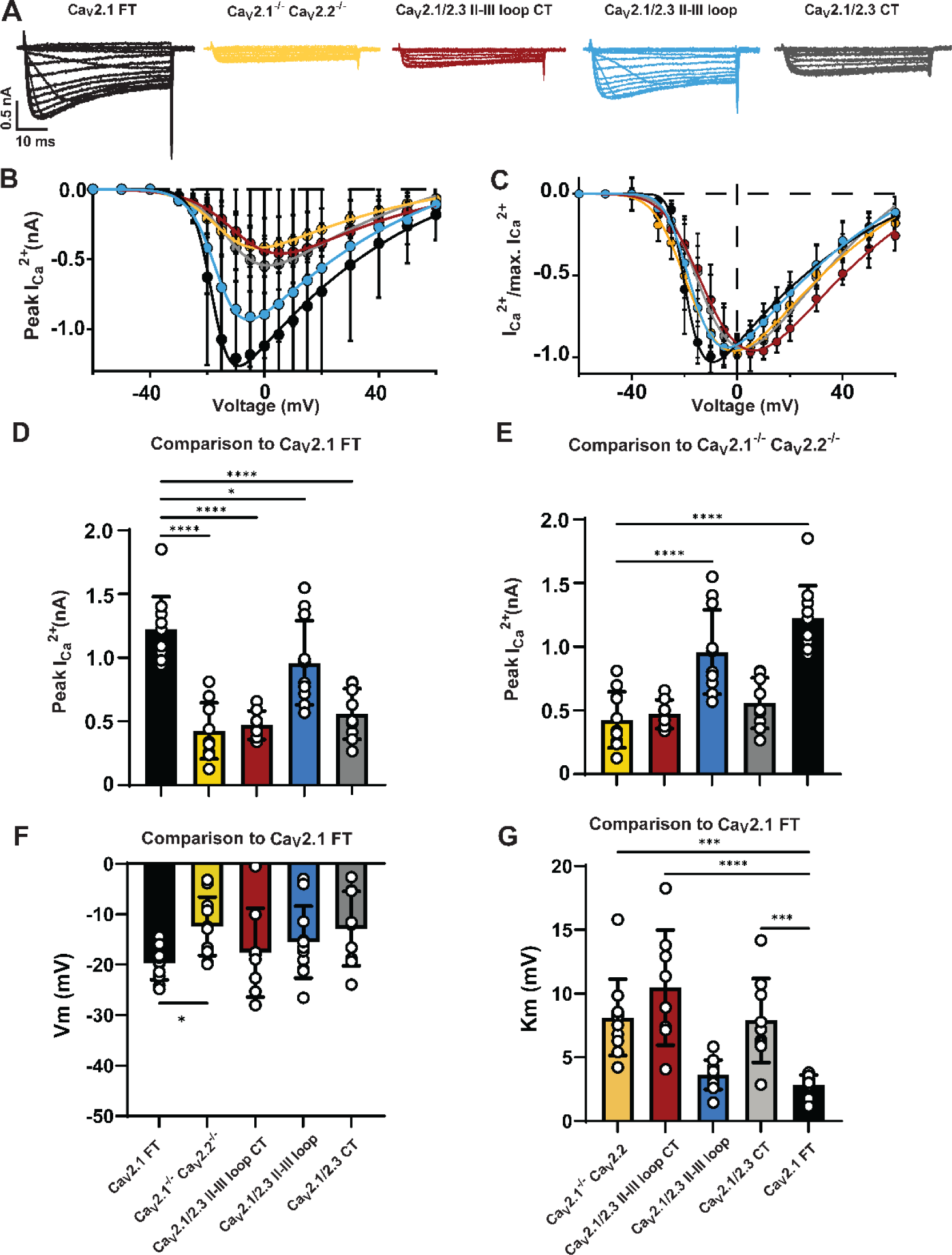
The Ca_V_2.1/2.3 α_1_ CT chimera cannot rescue Ca^2+^ currents to Ca_V_2.1 FT levels, however the Ca_V_2.1/2.3 loop II-III chimera partially rescues Ca^2+^ currents. Sample traces of Ca^2+^ current triggered by 50 ms voltage steps from -60 mV to -30 mV in 10 mV steps, -25 mV to +20 mV in 5 mV steps, and +30 mV to +60 mV in 10 mV steps. Black: calyces expressing Ca_V_2.1 FT (n=11). Yellow: Ca_V_2.1^-/-^/ Ca_V_2.2^-/-^ calyces (n=11). Red: calyces expressing Ca_V_2.1/2.3 II-III loop CT chimeric construct (n=8). Blue: calyces expressing Ca_V_2.1/2.3 II-III loop chimeric construct (n=11). Gray: calyces expressing Ca_V_2.1/2.3 CT chimeric construct (n=9). (B-C) IV relationship of Ca^2+^ currents (B) and normalized Ca^2+^ currents by maximal Ca^2+^ currents (I/I_max_; C) plotted against voltage steps. (D-E) Bar graph shows absolute peak Ca^2+^ current amplitudes, one-way ANOVA, Dunnett’s test, Ca_V_2.1 FT is used as positive control (D) and Ca_V_2.1^-/-^/ Ca_V_2.2^-/-^ is used as negative control (E). (F-G) Bar graph shows voltage dependent activation kinetics of Ca_V_2 chimeras. Half-maximal activation voltage (V*_m_*) (F) and voltage-dependence of activation (K*_m_*) (G) is plotted. One-way ANOVA, Dunnett’s test, Ca_V_2.1 FT is used as positive control.

Subsequently, we expressed the Ca_V_2.1/2.3 chimeras and performed direct presynaptic Ca^2+^ recordings. All chimera constructs were compared to Ca_V_2.1 α_1_ subunit FT and Ca_V_2.1^-/-^/ Ca_V_2.2^-/-^ calyces to determine rescue efficiency (Fig. 3). Since we did not find any differences in the calyx of Held size with expression of the Ca_V_2 chimeras (Table 3), we reported data as maximal Ca^2+^ currents amplitude at all voltages. Current to voltage (IV) analysis revealed significant reduction in peak Ca^2+^ current amplitudes with all three chimeras compared to Ca_V_2.1 FT (Fig. 3) However, only expression of the Ca_V_2.1/2.3 loop II-III led to maximum Ca^2+^ currents that were larger than that of the Ca_V_2.1^-/-^/ Ca_V_2.2^-/-^ (Maximum Ca^2+^ current amplitudes (pA) Ca_V_2.1^-/-^/ Cav2.2^-/-^: -425.5 ± 219.5 (n=11) vs. Ca_V_2.1/2.3 II-III loop: -958.1 ± 331.3 (n=11), p<0.0001) (Fig. 3E).

**Table3.** Capacitances of calyx of Held expressing CaV2 hybrid channels.

| Parameter | Mean | SD | n | P | statistics |
| --- | --- | --- | --- | --- | --- |
| **Membrane capacitance C<sub>slow</sub> (pF)** |  |  |  |  |  |
| Wildtype (control) | 20.59 | 4.220 | 10 | p=0.9246 (n.s.) | One-Way ANOVA<br>Dunnett's Test |
| Cav2.1 FT | 23.80 | 5.150 | 11 | control |  |
| Cav2.1<sup>-/-</sup>/ Cav2.2<sup>-/-</sup> | 22.85 | 8.831 | 11 | p=0.9996 (n.s.) |  |
| Cav2.1/2.3 II-III loop CT | 21.72 | 6.096 | 8 | p=0.9990 (n.s.) |  |
| Cav2.1/2.3 II-III loop | 25.76 | 6.045 | 11 | p=0.9960 (n.s.) |  |
| Cav2.1/2.3 CT | 19.06 | 4.894 | 9 | p=0.6135 (n.s.) |  |
| Cav2.1 Δ2016 | 27.26 | 3.420 | 6 | p=0.9522 (n.s.) |  |
| Cav2.1/2.3 CT 2016-2368 | 28.22 | 7.622 | 12 | p=0.6103 (n.s.) |  |
| Cav2.1/2.3 CT 1967-2368 | 27.47 | 8.513 | 9 | p=0.8646 (n.s.) |  |
| Cav2.1/2.2 CT | 21.94 | 9.095 | 11 | p=0.9991 (n.s.) |  |
| Pun Cav2.1 FT | 21.44 | 5.828 | 7 | p=0.9955 (n.s.) |  |
| Pun Cav2.1/2.2 CT | 23.94 | 4.943 | 9 | p>0.9999 (n.s.) |  |
| Pun Cav2.1/2.3 CT | 26.04 | 7.309 | 10 | p=0.9916 (n.s.) |  |

Although we have demonstrated that changes in presynaptic Ca^2+^ currents correlate with changes in channel numbers (Lubbert *et al*., 2017; Dong *et al*., 2018; Lubbert *et al*., 2019a), and presynaptic Ca_V_2.3 levels are significantly less than Ca_V_2.1 and correspond to limited contribution of Ca_V_2.3 channels to the presynaptic Ca_V_2 current subtype contribution (Wu *et al*., 1999) it was possible that the reduction in presynaptic currents with the chimeric constructs could be due to changes in the channel open probability (Po) or general mechanisms that regulate Ca_V_2 channel numbers in the membrane that is not specific to regulation of presynaptic abundance. To rule out these possibilities, we expressed our Ca_V_2.1/2.3 chimeras in HEK293T cells and measured the gating current (I*_gating_*) and peak tail current (I*_tail_*) at the reversal potential for Ca_V_2 channels (Fig 4). The time integral of I_gating_ (Q_max_) is an estimate of channel number (N). Analysis of Q_max_ found no difference between the chimeras, indicating that there was no change in N between the chimeras. Subsequently we plotted peak I*_tail_* vs Q_max_ and performed linear regression for the constructs to determine the index of the relative Po for each construct (Fig 4). Results of our analysis revealed comparable slopes between linear regressions of the individual constructs indicating that the relative Po of the Ca_V_2.1/2.3 chimeric channels was similar.

**Figure 4.**
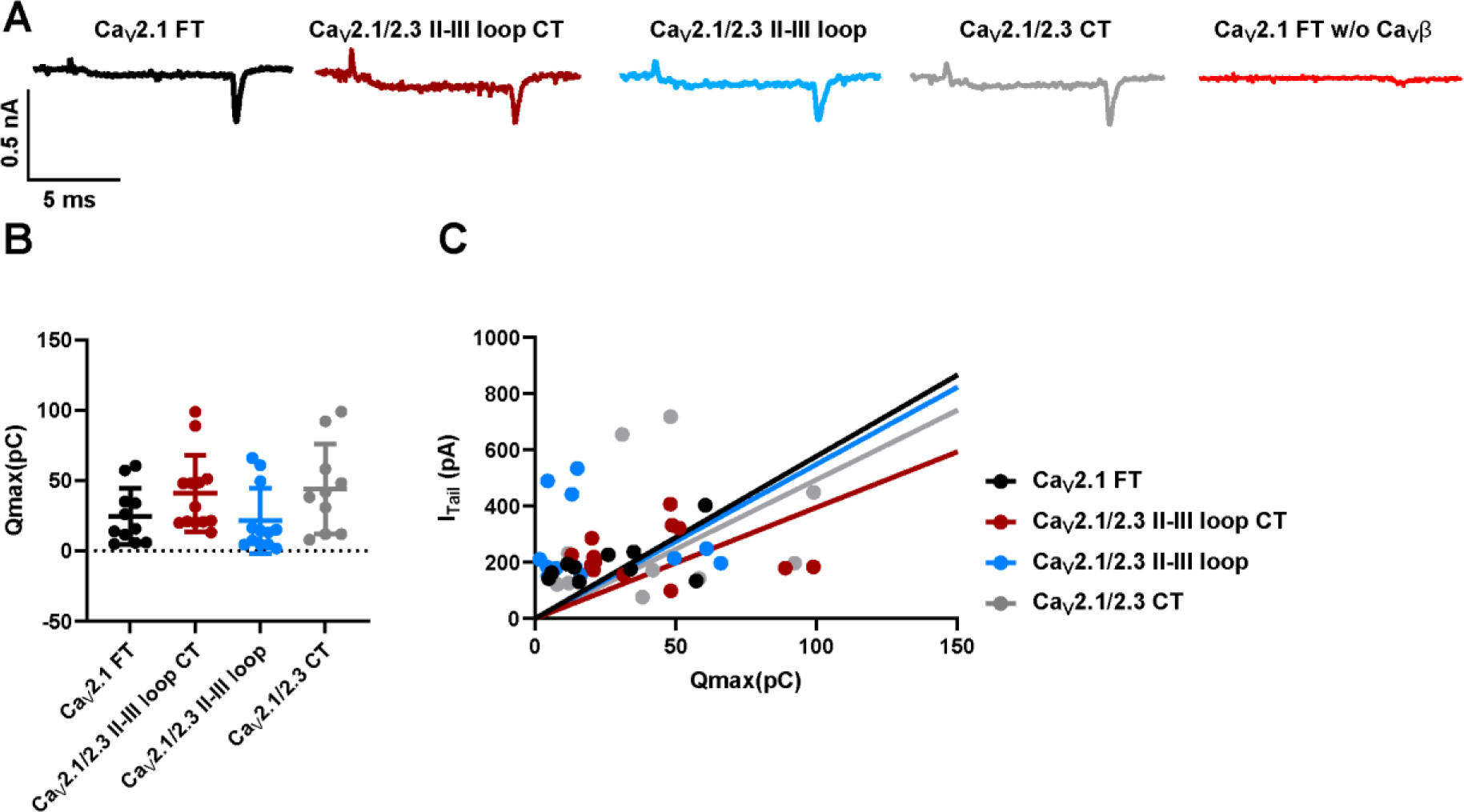
Ca_V_2 chimeras have similar expression level and open probability in HEK293T cells. (A) CMV promoter is used to drive Ca_V_2 chimera expressions in HEK293T cells. Example Ca^2+^ current traces to 10ms depolarization from -80mV holding potential to the reverse potential +50mV for HEK293T cells transfected with CMV Ca_V_2.1 FT (black, n=11), CMV Ca_V_2.1/2.3 II-III loop CT (dark red, n=13), CMV Ca_V_2.1/2.3 II-III loop (blue, n=12), CMV Ca_V_2.1/2.3 CT (gray, n=10) and CMV Ca_V_2.1 FT without CMV Ca_V_β4 co-expression (red, n=7). (B-C) Bar graph shows Q*_max_* (B) and linear regression plots (C) shows ratio between I_tail_ and Q_max_ in HEK293T cells co-transfected with CMV Ca_V_2 chimeras and CMV Ca_V_β4.

Based on these results, we demonstrate that Ca_V_2.1 loop II-III and Ca_V_2.1 CT are positive regulators of presynaptic Ca_V_2.1 abundance but differentially regulate presynaptic Ca_V_2.1 abundance.

### The Ca_V_2.1 α_1_ subunit loop II-III and Ca_V_2.1 α_1_ subunit CT do not independently control Ca_V_2.1 preference

Ablation of Ca_V_2 subtypes leads to increased presynaptic levels of other Ca_V_2 subtypes (Inchauspe *et al*., 2004; Lubbert *et al*., 2017) (Fig 1D, F), and overexpression of Ca_V_2.1 results in increased presynaptic Ca_V_2.1 levels and competes away Ca_V_2.2, but not vice versa (Lubbert *et al*., 2019a). Therefore, it is postulated that presynaptic Ca_V_2 abundance is correlated to Ca_V_2 subtype preference. Although we found a reduction of presynaptic Ca^2+^ current with the Ca_V_2.1/2.3 chimeras, whether these regions regulated Ca_V_2.1 preference or just abundance is unknown. Therefore, it was important to elucidate if motifs that regulate presynaptic Ca_V_2 subtype abundance are also utilized to determine Ca_V_2 subtype preference or if they were distinct. The remaining Ca^2+^ currents from calyces expressing the Ca_V_2.1/2.3 chimeras could be mediated by: 1) a mixture of Ca_V_2.1/2.3 chimeras and Ca_V_2.3 channels, 2) solely Ca_V_2.1/2.3 chimeras, or 3) solely Ca_V_2.3 channels. Since the presynaptic Ca^2+^ currents in the Ca_V_2.1^-/-^ /Ca_V_2.2^-/-^ calyx of Held were completely Aga insensitive, and the Ca_V_2.1/2.3 chimeras still maintain Aga binding sites and are the Aga sensitive, we measured Aga sensitivity of Ca^2+^ currents with the Ca_V_2.1/2.3 chimeras to distinguish between the different possibilities. In scenario 1, the Ca_V_2.1/2.3 chimeras are localized to the presynaptic membrane but do not completely away or only partially compete away Ca_V_2.3 channels and would result in a mix of Aga sensitive and insensitive Ca^2+^ currents. In scenario 2, Ca_V_2.1/2.3 chimeras are localized to the presynaptic membrane and compete away Ca_V_2.3 channels and would result in similar Aga sensitivity as the Ca_V_2.1 FT. In scenario 3, Ca_V_2.1/2.3 chimeras are not localized to the presynaptic membrane with only Ca_V_2.3 currents remaining and would result in similar Aga insensitivity as the Ca_V_2.1^-/-^/ Ca_V_2.2^-/-^ calyx of Held. Analysis of Aga sensitivity of Ca_V_2.1/2.3 II-III or Ca_V_2.1/2.3 CT showed no difference compared to the Ca_V_2.1 FT (Fig. 5). However, we found partial Aga sensitivity with Ca_V_2.1/2.3 II-III+ CT (Aga sensitive fraction (%): Ca_V_2.1 FT 90.43 ± 5.97 (n=7) vs. Ca_V_2.1/2.3 II-III+ CT 69.67 ± 12.93 (n=9), p=0.0017) (Fig. 5). Based on our data, we demonstrate that both Ca_V_2.1/2.3 chimeras with the loop II-III or CT swap are able to compete away Ca_V_2.3 channels (Fig. 5). Therefore, we conclude that Ca_V_2.1 loop II-III and the CT do not independently control Ca_V_2.1 preference and motifs that regulate presynaptic Ca_V_2.1 preference are distinct from those that regulate presynaptic Ca_V_2.1 abundance.

**Figure 5.**
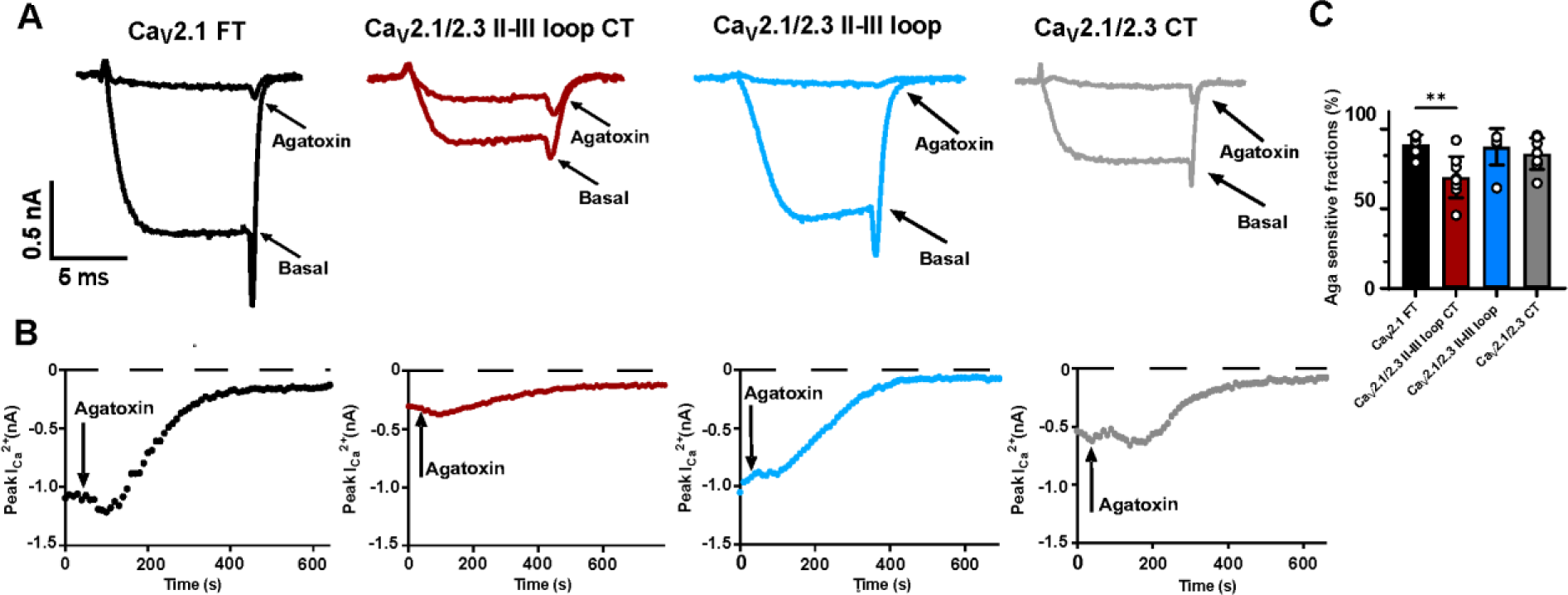
The Ca_V_2.1/2.3 α_1_subunit chimeras compete away Ca_V_2.3 channel in the calyx of Held. (A) Agatoxin isolation of presynaptic Ca_V_2.1/2.3 chimeras in calyces expressing Ca_V_2.1 FT (black, n=7), Ca_V_2.1/2.3 II-III loop CT (red, n=9), Ca_V_2.1/2.3 II-III loop (blue, n=7), and Ca_V_2.1/2.3 CT (gray, n=8). Basal and reduced Ca^2+^ current traces after applying 200 nM ω-Aga IVA are labelled. (B) Reduction of Agatoxin sensitive Ca^2+^ current over time. Ca^2+^ current amplitudes are plotted against time, 200 nM ω-Aga IVA was applied at 5^th^ pulse (50s). (C) Bar graph shows the ω-Aga IVA sensitive reduction fraction in calyces expressing Ca_V_2.1 FT, Ca_V_2.1/2.3 II-III loop CT, Ca_V_2.1/2.3 II-III loop, and Ca_V_2.1/2.3 CT.

### The Ca_V_2.3 α_1_ subunit CT negatively regulates presynaptic Ca_V_2.1 abundance while the Ca_V_2.1 CT aa. 1933-1966 positively regulates presynaptic Ca_V_2.1 abundance

The Ca_V_2.3 CT contains a Calmodulin binding domain (CBD) but does not contain many motifs postulated to mediate presynaptic Ca_V_2.1 subtype abundance (Kamp *et al*., 2012; Nanou & Catterall, 2018). In addition, proposed Ca_V_2.1 CT motifs (aa 2016 onwards) are not essential determinants of presynaptic Ca_V_2.1 abundance (Lubbert *et al*., 2017). Thus, we hypothesized that the 1) Ca_V_2.3 CT acts as a negative regulator of presynaptic Ca_V_2 abundance, and/or 2) the Ca_V_2.1 CBD regulates presynaptic Ca_V_2 abundance. Therefore, to elucidate the mechanism of action of the Ca_V_2.3 CT, we created two additional Ca_V_2.1/2.3 CT chimeras. They were the <u>Ca_V_2.1/2.3 CT 1967-2368</u>, which replaces the Ca_V_2.1 CT with the Ca_V_2.3 CT from Ca_V_2.1 aa. 1967 onwards and contains the Ca_V_2.3 CBD, and the <u>Ca_V_2.1/2.3 CT 2016-2368</u> which replaces the Ca_V_2.1 CT with the Ca_V_2.3 CT from Ca_V_2.1 aa 2016 onwards (Fig. 6A). Subsequently, we expressed these chimeras and Ca_V_2.1 Δ2016, which deletes all motifs, AZ protein binding sites, and the secondary Ca_V_β interaction site(Lubbert *et al*., 2017) and measured their maximum Ca^2+^ amplitudes to determine rescue efficiency (Fig. 6). As previously published. we found no difference in maximum Ca^2+^ currents between Ca_V_2.1 Δ2016 and Ca_V_2.1 FT (Fig. 6B-C), thereby confirming that motifs in the Ca_V_2.1 CT aa. 2016 onwards are not essential for controlling presynaptic Ca_V_2.1 abundance. In contrast, expression of Ca_V_2.1/2.3 CT 2016-2368 and Ca_V_2.1/2.3 CT 1967-2368 resulted in a reduction in the maximum Ca^2+^ current amplitudes (Ca_V_2.1 Δ2016: -1241 ± 207.4 (n=6) vs. Ca_V_2.1/2.3 CT 2016-2368: -934.1 ± 210.6 (pA) (n=12), p=0.0209; Ca_V_2.1 Δ2016: -1241 ± 207.4 (n=6) vs. Ca_V_2.1/2.3 CT 1967-2368: -839.5 ± 251.9 (pA) (n=8), p=0.0047) (Fig. 6D-F), but the maximum Ca^2+^ current amplitudes with either Ca_V_2.1/2.3 CT 1967-2368 or Ca_V_2.1/2.3 CT 2016-2368 were larger than the Ca_V_2.1/2.3 CT 1933-2368 ((pA) Ca_V_2.1/2.3 CT aa. 1933-2368: -557.7 ± 198.9 (n=9) vs. Ca_V_2.1/2.3 CT 2016-2368: -934.1 ± 210.6 (n=12), p=0.0013; Ca_V_2.1/2.3 CT aa. 1933-2368: -557.7 ± 198.9 (n=9) vs. Ca_V_2.1/2.3 CT 1967-2368: -839.5 ± 251.9 (n=8), p=0.0321) (Fig. 6D-F). Therefore, we conclude that the Ca_V_2.3 CT aa. 1952-2273 contains a motif or motifs that negatively regulate presynaptic Ca_V_2.3 abundance and the Ca_V_2.1 CBD is not a positive regulator of presynaptic Ca_V_2.1 abundance. Furthermore, we identified the region between Ca_V_2.1 CT aa. 1933-1966 positively regulates presynaptic Ca_V_2.1 abundance.

**Figure 6.**
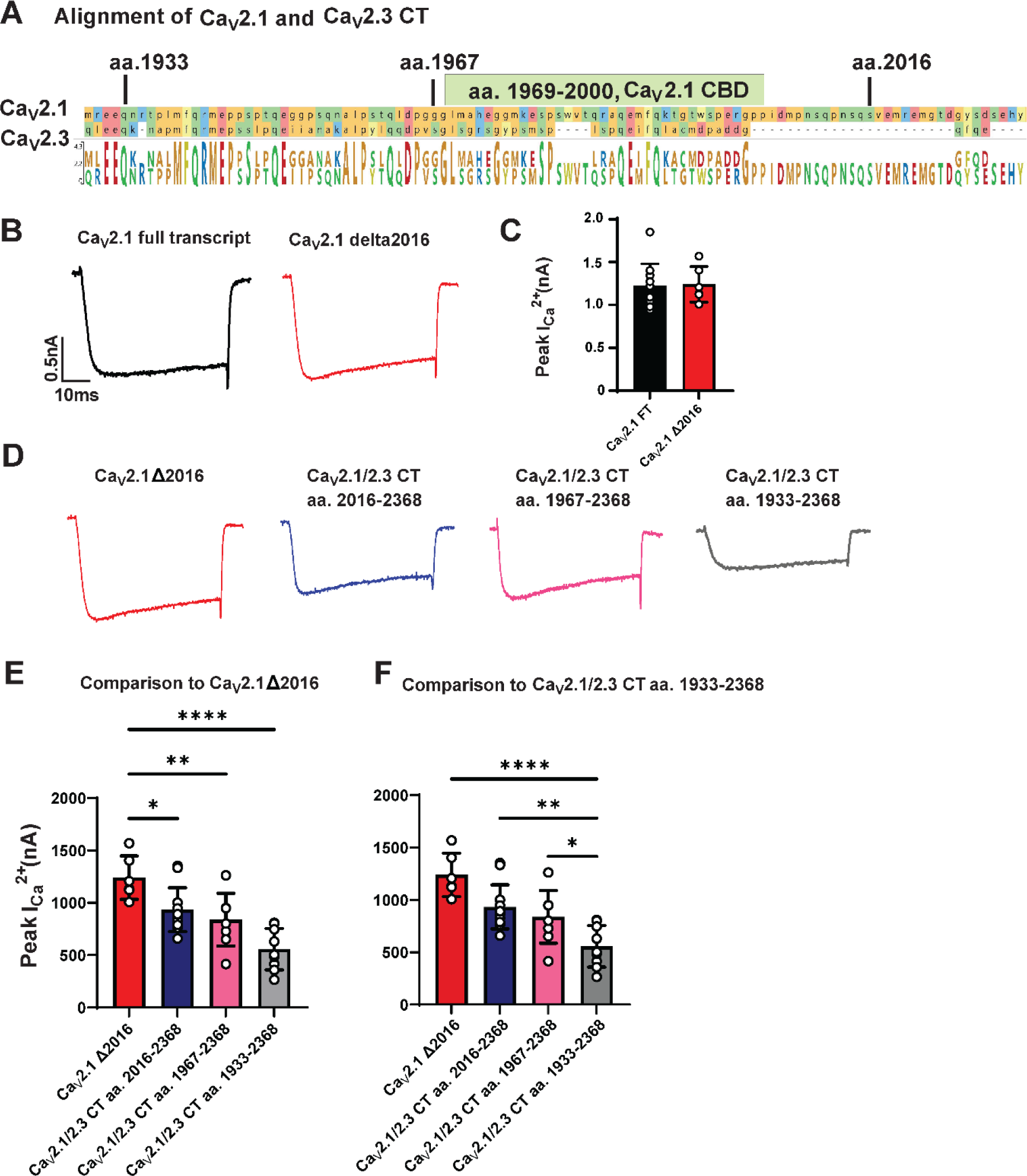
The Ca_V_2.3 CT negatively regulates presynaptic Ca_V_2 abundance, while the Ca_V_2.1 CT region aa. 1933-1966 promotes Ca_V_2.1 abundance. (A) Amino acid alignment of Ca_V_2.1 and Ca_V_2.3 CT. Sequences on top and bottom represent Ca_V_2.1 CT and Ca_V_2.3 CT, respectively. Amino acid numbers of swapped sites in three Ca_V_2.1/2.3 chimeras are indicated in black. Ca_V_2.1 CBD is highlighted in green. (B) Example Ca^2+^ current traces to 50 ms depolarization from -80mV holding potential to the maximal activation voltage step (-15mV to +15mV, 5 mV steps) for Ca_V_2.1 FT (black, n=11) and Ca_V_2.1 Δ2016 (red, n=6). (C) Bar graph shows absolute peak Ca^2+^ currents amplitude between Ca_V_2.1 CT and Ca_V_2.1 Δ2016, unpaired t test. (D) Example Ca^2+^ current traces to 50ms depolarization from -80mV holding potential to the maximal activation voltage step (-15mV to +15mV, 5 mV steps) for Ca_V_2.1 Δ2016, Ca_V_2.1/2.3 CT aa. 2016-2368 (n=12); Ca_V_2.1/2.3 CT 1967-2368 (n=8) and Ca_V_2.1/2.3 CT 1933-2368 (n=9). (E-F) Bar graph shows peak Ca^2+^ currents amplitude, one-way ANOVA, Dunnett’s test, Ca_V_2.1 Δ2016 is used as positive control (E) and Ca_V_2.1/2.3 CT aa. 1933-2368 is used as negative control (F).

### The Ca_V_2.2 CT does not control presynaptic Ca_V_2.1 abundance

Since we identified that the Ca_V_2.3 CT was a negative regulator of presynaptic Ca_V_2.3 abundance, we set out to determine if this was unique to the Ca_V_2.3 CT. Thus, to address if Ca_V_2.2 CT negatively regulated presynaptic Ca_V_2 abundance, we created a Ca_V_2.1/2.2 CT chimera and cloned this into our HdAd vectors. Western blot analysis demonstrated no issues with expression of the Ca_V_2.1/2.2 CT chimera (Fig. 7A). Subsequently we expressed the Ca_V_2.1/2.2 CT chimera and performed IV analysis on presynaptic Ca^2+^ currents. Results of our IV analysis revealed that there was no difference in the maximum Ca^2+^ current between the Ca_V_2.1/2.2 CT chimera and Ca_V_2.1 FT (Fig. 7B-E). However, compared to Ca_V_2.1 FT, there appeared to be unequal variances of maximum Ca^2+^ current amplitudes with Ca_V_2.1/2.2 CT that was marginally significant (F-test comparing variances p=0.0607) (Fig. 7B-E). Therefore, our data suggests that the Ca_V_2.2 CT is not a negative regulator of presynaptic abundance.

**Figure 7.**
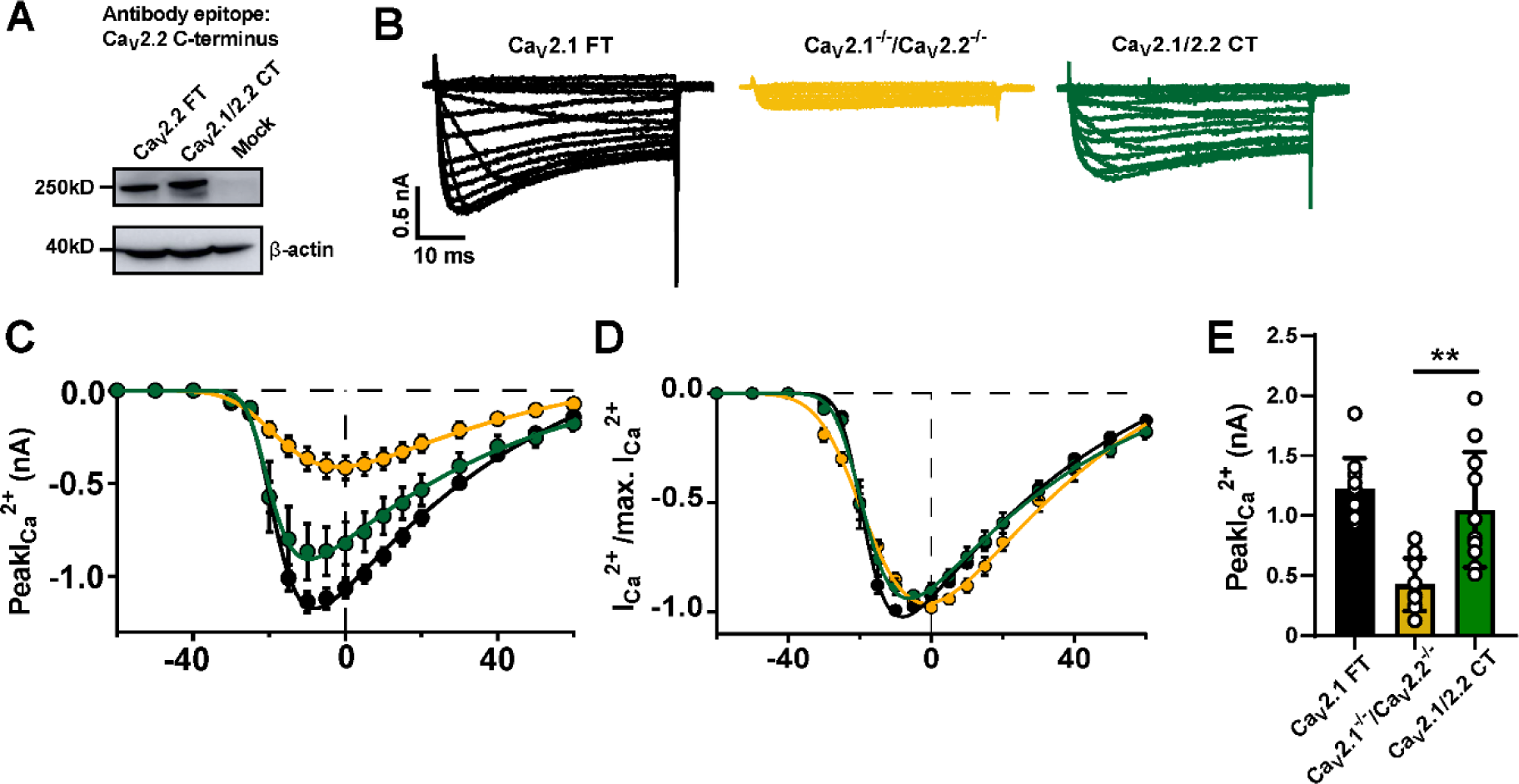
Ca_V_2.1/2.2 CT chimera can be efficiently expressed and rescues Ca^2+^ currents to Ca_V_2.1 FT level with high variability. (A) Western Blot validated protein expression of Ca_V_2.1/2.2 CT. HEK293 cells are infected by HdAd Ca_V_2.1/2.2 CT and Ca_V_2.2 FT. Antibody targeting Ca_V_2.2 CT shows the protein expression of Ca_V_2.2 FT and Ca_V_2.1/2.2 CT. Mock, negative control, protein lysis from uninfected HEK293 cells. (B) Exemplary Ca^2+^ currents triggered by 50 ms voltage steps from - 60 mV to -30 mV in 10 mV steps, -25 mV to +20 mV in 5 mV steps, and +30 mV to +60 mV in 10 mV steps. Black: calyces expressing Ca_V_2.1 FT (n=11). Yellow: Ca_V_2.1^-/-^/ Ca_V_2.2^-/-^ calyces (n=11). Green: calyces expressing Ca_V_2.1/2.2 CT (n=11). (C-D) IV relationship of Ca^2+^ currents (C) and normalized Ca^2+^ currents by maximal Ca^2+^ currents (I/I_max_; D) plotted against voltage steps. (E) Bar graph shows absolute peak Ca^2+^ current amplitudes. One-way ANOVA, Dunnett’s test, Ca_V_2.1 FT is used as positive control, Ca_V_2.1^-/-^/ Ca_V_2.2^-/-^ is used as negative control.

### High level overexpression of Ca_V_2.1/2.2 CT rescues presynaptic Ca^2+^ currents while Ca_V_2.1/2.3 CT does not

High level overexpression of Ca_V_2.1 α_1_ subunit results in increased presynaptic Ca_V_2.1 levels and compete away Ca_V_2.2 and Ca_V_2.3 channels, however high-level overexpression of Ca_V_2.2 α_1_ subunit does not result in a reduction of Ca_V_2.1 levels at the calyx of Held (Lubbert *et al*., 2019a). This suggests that upstream regulatory steps, such as protein trafficking and membrane insertion steps, are integral to Ca_V_2.1 level preference and abundance (Ferron *et al*., 2021; Young & Veeraraghavan, 2021). It is known that high levels of overexpression can saturate regulatory steps that control trafficking (Lubbert *et al*., 2019a). Therefore, to determine if these regulatory step(s) could be saturated, we used the high-level overexpression Pun cassette (Young & Neher, 2009; Montesinos *et al*., 2011; Lubbert *et al*., 2019a) to drive expression of Ca_V_2.1/2.2 CT and Ca_V_2.1/2.3 CT chimeras and determine if we could increase the presynaptic Ca_V_2 chimera current levels and compared their maximum Ca^2+^ current amplitudes to Pun Ca_V_2.1 FT. We found no difference in presynaptic Ca^2+^ currents with Pun Ca_V_2.1/2.2 CT compared to Pun Ca_V_2.1 FT (Fig. 8). However, maximum Ca^2+^ current amplitudes with Pun Ca_V_2.1/2.3 CT were lower than Pun Ca_V_2.1 FT (Pun Ca_V_2.1 FT: -1438 ± 198.9 (n=7) vs. Pun Ca_V_2.1/2.3 CT: -772.8 ± 291.8 (n=10), (pA), p=0.0006) (Fig. 8). Moreover, the maximum Ca^2+^ current amplitudes with Pun Ca_V_2.1/2.3 CT showed no significant difference than syn Ca_V_2.1/2.3 CT (Fig. 8). Therefore, we demonstrate that the Ca_V_2.2 CT is not a negative regulator of presynaptic Ca_V_2.2 abundance and the Ca_V_2.1/2.2 CT chimera can be forced to increase expression level, however, the negative regulator motif in Ca_V_2.3 CT restrains the upstream regulatory steps which results in low abundance of Ca_V_2.3 channels in the presynaptic terminal.

**Figure 8.**
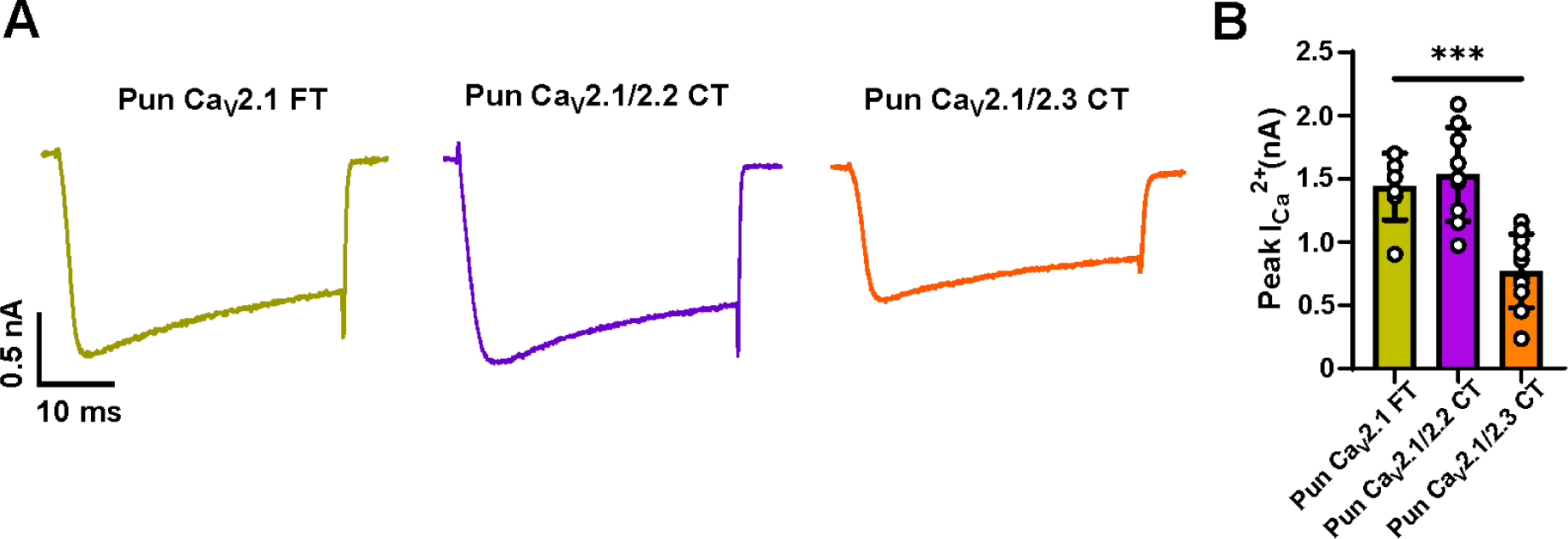
High level overexpression of the Ca_V_2.1/2.3 α_1_ CT cannot rescue Ca^2+^ currents to Pun Ca_V_2.1 levels while Pun Ca_V_2.1/2.2 α_1_ CT can. (A) High-level overexpression cassette pUNISHER (Pun) is used to drive Ca_V_2.1 FT, Ca_V_2.1/2.2 CT and Ca_V_2.1/2.3 CT chimeric constructs. Example Ca^2+^ current traces to 50ms depolarization from -80mV holding potential to the maximal activation voltage step (-15mV to +15mV, 5 mV steps) for calyces expressing Pun Ca_V_2.1 FT (green, n=7), Pun Ca_V_2.1/2.2 CT (purple, n=9) and Pun Ca_V_2.1/2.3 CT (orange, n=10). (B) Bar graph shows averaged peak Ca^2+^ currents for Pun Ca_V_2.1 FT, Pun Ca_V_2.1/2.2 CT and Pun Ca_V_2.1/2.3 CT constructs. One-way ANOVA, Dunnett’s test, Pun Ca_V_2.1 FT is used as positive control.

## Discussion

In this study, we provide insight into how presynaptic Ca_V_2 subtype abundance is determined by the Ca_V_2 α_1_ subunit. Based on our data, we demonstrate that the cytoplasmic loop II-III and CT regions of the Ca_V_2.1 α_1_ subunit are positive regulators of presynaptic Ca_V_2 abundance, while Ca_V_2.3 CT negatively regulates presynaptic Ca_V_2 abundance. In addition, we find that Ca_V_2.2 CT is not a key determinant of presynaptic Ca_V_2 abundance. Furthermore, we demonstrate that Ca_V_2.1 α_1_ subunit cytoplasmic loop II-III and CT regions do not independently control presynaptic Ca_V_2.1 preference. Therefore, we propose that the motifs controlling presynaptic Ca_V_2.1 preference are distinct from those regulating Ca_V_2.1 levels and they may act synergistically in pathways that regulate presynaptic Ca_V_2.1 preference and levels.

### The Ca_V_2 α_1_ subunit II-III loop regulation of presynaptic Ca_V_2.1 abundance

By swapping the loop II-III domain between Ca_V_2.1 and Ca_V_2.3 we demonstrated that the Ca_V_2.1 loop II-III region positively regulates presynaptic Ca_V_2.1 abundance but does not regulate preference. All Ca_V_2 subunit loop II-III domains contain a synprint region which is the site of interaction between the SNARE proteins syntaxin and SNAP-25 (Zhong *et al*., 1999; Mochida *et al*., 2003; Yokoyama *et al*., 2005). However, the proposed syntaxin binding domain found in Ca_V_2.1 and Ca_V_2.2 is not found in the Ca_V_2.3 α_1_ subunit synprint region (Kamp *et al*., 2005; Yokoyama *et al*., 2005; Rajapaksha *et al*., 2008). Therefore, our findings that the Ca_V_2.1/2.3 loop II-III chimera resulted in presynaptic Ca^2+^ current reductions indicates that potential differences in syntaxin binding between the different subtypes of Ca_V_2 α_1_ subunits may regulate presynaptic Ca_V_2 abundance. Studies from cultured hippocampal neurons, Ca_V_2.1 dominant presynaptic terminals, found that partial deletions in the Ca_V_2.2 synprint region that impact SNARE binding resulted in less efficient Ca_V_2.2 targeting to presynaptic terminal (Szabo *et al*., 2006). Similar partial deletions in the Ca_V_2.1 synprint region reduced Ca_V_2.1 localization to release sites in cultured superior cervical ganglion (SCG) neurons which utilize Ca_V_2.2 dominant presynaptic terminals (Mochida *et al*., 2003). In addition, splice variants of Ca_V_2.1 lacking the synprint region reduce incorporation into neuroendocrine cells (Rajapaksha *et al*., 2008). Since partial deletions of Ca_V_2.2 synprint do not impair axonal targeting(Szabo *et al*., 2006), it is possible that the Ca_V_2.3 synprint region does not regulate axonal trafficking but rather regulates incorporation into the presynaptic terminal. Although the Ca_V_2.3 synprint region chimera resulted in reduced presynaptic Ca_V_2 chimera levels, there was no change in the proportion of Agatoxin sensitive currents. Therefore, this indicates the mechanisms that control Ca_V_2.1 preference are not determined by the loop II-III region binding interactions.

An alternative interpretation of our data is that the loop II-III swaps resulted in changes in increased inactivation of the Ca_V_2.1/2.3 channels which resulted in a reduction in presynaptic Ca^2+^ currents that was unrelated to abundance. However, deletion of the entire synprint site which contains syntaxin and SNAP25 binding sites from the Ca_V_2.1 α_1_ subunit has similar activation and a positive shift in the inactivation curves and there was no difference in normalized conductance at -80 mV and -70 mV (Mochida *et al*., 2003). Furthermore, Ca_V_2.3 channels have a half inactivation voltage(Pereverzev *et al*., 2002) that is similar to Ca_V_2.1 (Mochida *et al*., 2003). More importantly, we demonstrated the relative *Po* and N between the Ca_V_2.1/2.3 loop II-III channels and Ca_V_2.1 channels were similar in HEK293T cells.

### The Ca_V_2 α_1_ subunit CT differentially regulates presynaptic Ca_V_2 subtype abundance

We found that the Ca_V_2.2 CT and Ca_V_2.3 CT had differential effects on presynaptic Ca_V_2 abundance but did not impact Ca_V_2 preference. Our data demonstrated that Ca_V_2.1/2.3 CT chimera results in a severe reduction in presynaptic Ca^2+^ currents with no difference in the Po or N. Therefore, our data suggests that the Ca_V_2.3 CT is a negative regulator of presynaptic Ca_V_2.3 abundance. Prior studies (Lubbert *et al*., 2017) and our findings here revealed that the Ca_V_2.1 CT Δ2016, which deletes multiple AZ protein binding sites and other motifs restores presynaptic Ca^2+^ current levels to Ca_V_2.1 FT levels. These data demonstrate these interactions and motifs are not essential for controlling presynaptic Ca_V_2.1 abundance. However, replacing the Ca_V_2.1 CT from aa. 2016 onwards with corresponding Ca_V_2.3 CT aa. 1952-2273 resulted in a reduction of presynaptic Ca^2+^ currents compared to Ca_V_2.1 CT Δ2016. Furthermore, overexpression of Ca_V_2.1/2.3 CT chimera by pUNISHER cassette was similar to presynaptic Ca^2+^ currents levels with the low-level synapsin expression cassette. Although it is possible that the Ca_V_2.3 CT contains motif that alters tonic modulation of the channels, Ca_V_2.3 channels have a half inactivation voltage (Pereverzev *et al*., 2002) that is similar to Ca_V_2.1 (Mochida *et al*., 2003). Finally, Ca_V_2.3 levels are significantly less than Ca_V_2.1 and correspond to limited contribution of Ca_V_2.3 channels to the presynaptic Ca_V_2 current subtype contribution (Wu *et al*., 1999).

Therefore, we propose that Ca_V_2.3 CT aa. 1952-2273 contains a motif that prevents presynaptic Ca_V_2 targeting or incorporation into the plasma membrane. In addition, our results indicate that CT dependent mechanism of regulation of presynaptic Ca_V_2 abundance is dominant to the loop II-III region mechanisms of action as Ca_V_2.1/2.3 CT chimera resulted in significantly lower presynaptic Ca^2+^ currents than those with the Ca_V_2.1/2.3 loop II-III.

Calmodulin (CaM) is considered a core component of Ca_V_2 channel complexes and interacts with the Cav2.1 channels through a bipartite interaction with the calmodulin binding domain (CBD) and IQ-like motif (IM) domain to regulate calcium dependent facilitation (CDF) and inactivation (CDI) (Lee *et al*., 2003; Chaudhuri *et al*., 2004; Mori *et al*., 2008). However, how CaM interactions regulate presynaptic Ca_V_2.1 levels is unclear. The Ca_V_2.1/2.3 CT aa. 1967-2368 chimera, which replaces the Ca_V_2.1 CBD with the Ca_V_2.3 CBD, showed no difference in rescuing efficiency compared to Ca_V_2.1/2.3 CT aa. 2016-2368 chimeras. However, we found that presynaptic Ca^2+^ currents with Ca_V_2.1/2.3 CT aa. 1967-2368 chimera were larger than that with Ca_V_2.1/2.3 CT aa. 1933-2368 chimera. Therefore, our results demonstrated that the Ca_V_2.1 CBD does not control presynaptic Ca_V_2.1 abundance but the linker region of Ca_V_2.1 CT aa. 1933-1966 between the IQ domain and the CBD plays a role in regulation of Ca_V_2.1 levels. Since the CaM/CBD complex interacts with the IM domain (Nanou *et al*., 2016; Serra *et al*., 2018), it is possible that the Ca_V_2.3 CBD and linker region with Ca_V_2.1 IQ domain result in a different structural conformation of the CT that impacts presynaptic Ca_V_2 abundance. Interestingly, the combination of the mutation (IM-AA) in the IM domain and CBD deletion in SCG neurons results in an increased peak Ca^2+^ currents levels, while the IM-AA or CBD deletion alone did not alter Ca^2+^ current levels (Mochida *et al*., 2008). Therefore, the IQ domain and CBD may synergistically regulate Ca_V_2.1 level. However, the molecular mechanisms how these regions impact Ca_V_2.1 abundance is unknown and future experiments are needed.

In contrast to the Ca_V_2.1/2.3 CT chimera, we found no difference in the average presynaptic Ca^2+^ current levels with the Ca_V_2.1/2.2 CT chimera compared to Ca_V_2.1 FT. However, we observed a large variability of Ca^2+^ current amplitudes with Ca_V_2.1/2.2 CT that is marginally significant compared to Ca_V_2.1 FT rescue. Therefore, our data implies that the Ca_V_2.2 CT is not a key determinant of presynaptic Ca_V_2 abundance. Unlike the Ca_V_2.3 CT, high level overexpression of Ca_V_2.1/2.2 CT resulted in similar presynaptic Ca^2+^ current levels compared to high level overexpression of Ca_V_2.1 FT level and no change in variability. Therefore, we propose that the Ca_V_2.2 CT does not contain motifs that prevents presynaptic Ca_V_2.2 targeting or incorporation into the plasma membrane.

### Ca_V_2 subtype abundance and preference are regulated by distinct pathways

Although there were differences in the presynaptic Ca^2+^ current levels with the Ca_V_2.1/2.3 loop II-III and Ca_V_2.1/2.3 CT chimeras, they had similar Aga sensitivity as presynaptic Ca^2+^ currents with the Ca_V_2.1 FT. However, there was a significant Aga insensitive component of Ca^2+^ currents with Ca_V_2.1/2.3 loop II-III+ CT chimera despite having similar Ca^2+^ current levels as the Ca_V_2.1/2.3 CT and Ca_V_2.1^-/-^/ Ca_V_2.2^-/-^ background. Therefore, our data indicates that: 1) Presynaptic Ca_V_2 subtype abundance mechanisms are independent of those that control preference. 2) The loop II-III region and the CT region do not individually encode preference. 3) The Ca_V_2.1 II-III loop and CT region may act synergistically to control Ca_V_2 subtype preference. 4) Other motifs in the Ca_V_2.1 α_1_ subunit outside of the CT and synprint region are involved in regulating presynaptic Ca_V_2 subtype preference. Potential candidates that may control preference are the Ca_V_2 α_1_ subunit at AID in domain I-II which binds to the Ca_V_β subunit with high affinity(De Waard *et al*., 1994). Ca_V_β subunits are critical for regulating Ca_V_2 trafficking and Ca_V_β4 and Ca_V_β2a are located within the presynaptic terminal. Ca_V_2.1 channels have higher specificity for binding to Ca_V_β4 than Ca_V_2.2 and Ca_V_2.3 channels (Xie *et al*., 2007; Muller *et al*., 2010). Another possibility is that preference is not encoded by motifs in the cytoplasmic regions but is controlled by the Cacna2d protein family (Dolphin, 2012; Hoppa *et al*., 2012; Stephani *et al*., 2019). Knockout of α_2_δ3 subunit in spiral ganglion neurons results in reduced Ca_V_2.1 and Ca_V_2.3 components with no change in Ca_V_2.2 subtype levels (Stephani *et al*., 2019). In contrast overexpression of α_2_δ1 subunit decreases Ca_V_2.2 level in cultured hippocampal neurons (Pilch *et al*., 2022), but did not change other Ca_V_2 subtype levels. Furthermore, the globular bushy cells which give rise to the calyx of Held, only express *Cacna2d2* and *Cacna2d3*. In addition, other synapses that undergo developmental reduction of Ca_V_2.2 and Ca_V_2.3 express *Cacna2d1*. Finally, the α_2_δ binding preference does not correlate to presynaptic Ca_V_2 subtype levels (Voigt *et al*., 2016) while CACNA2D expression patterns do not correlate to presynaptic Cav2 subytpe levels (Cole *et al*., 2005). Therefore, given these conflicting findings more experiments will be needed to decipher if the Cacna2d proteins regulate preference.

While it is possible that the loop II-III domain and the CT domain act in concert to regulate presynaptic preference, it is also possible that they are not involved in preference mechanisms but rather combination of the loop II-III and CT domain swap construct dramatically impair the Ca_V_2 chimera trafficking to the presynaptic terminal. In this scenario, the Ca_V_2.1/2.3 II-III loop+ CT chimera does not effectively compete away native Ca_V_2.3 channels, which are normally restricted from the upstream pathways that regulate presynaptic trafficking. Therefore, native Ca_V_2.3 channels are also trafficked to the presynaptic terminals that express the Ca_V_2.1/2.3 II-III loop+ CT chimera. In this hypothetical scenario, the preference mechanism(s) has high affinity for the Ca_V_2.1/2.3 chimera channel and low affinity for the Ca_V_2.3 channels, however by law of mass action, Ca_V_2.3 channels now participate in molecular pathways that determine preference and results in the presence of presynaptic Ca_V_2.3 channels. Based on our model, this demonstrates how alternative splicing of the Ca_V_2 α_1_ subunits and potentially alternative splicing of molecules in pathways that control Ca_V_2 trafficking (Weiss & Zamponi, 2017), and insertion into the membrane may lead to developmentally changes in presynaptic Ca_V_2 subtype levels. However, the molecular machinery and the motifs that determine preference remain to be identified.

### Limitations of our study

By generating our Ca_V_2.1 chimeric constructs and expressing them at the calyx of Held we identified regions in the Ca_V_2 α_1_ subunit that control presynaptic Ca_V_2 abundances based on changes in presynaptic Ca^2+^ currents levels. However, although we did not directly measure N presynaptic Ca^2+^ currents directly correlates to changes in presynaptic Ca^2+^ channel abundance (Holderith *et al*., 2012; Nakamura *et al*., 2015; Nusser, 2018; Lubbert *et al*., 2019a; Rebola *et al*., 2019; Radulovic *et al*., 2020). Therefore, an alternate hypothesis is that measures of whole cell presynaptic Ca^2+^ currents do not reflect changes in presynaptic Ca^2+^ channel abundance but rather changes in intrinsic biophysical properties of the chimeric channels that result in reduced presynaptic Ca_V_2 channel activity, or differences in related to general mechanisms trafficking or translation. However, measurements in HEK293T cells showed no change in the peak currents between the Ca_V_2.1/2.3 chimeras and no change in the relative *Po* and N of the chimeric constructs in HEK293T cells. Moreover, all Ca_V_2 chimeras cDNA sequences were codon optimized and identical transgene cassettes were used to express these constructs. Therefore, reduced presynaptic Ca^2+^ currents are not due to changes of intrinsic biophysical properties, Po or translation levels. More importantly, this supports that the reductions in presynaptic Ca^2+^ currents reflect changes in presynaptic Ca_V_2 channel abundance. Finally, studies did not determine the exact molecular mechanism, such as channel trafficking, insertion, retention, that control presynaptic abundance but have identified specific motifs in the Ca_V_2 α_1_ subunit. Future studies will be needed to determine these steps. However, while presynaptic terminals appear to have a ceiling for total amount of calcium channels (Cao *et al*., 2004; Holderith *et al*., 2012; Nakamura *et al*., 2015; Nusser, 2018; Lubbert *et al*., 2019b; Rebola *et al*., 2019), specific slots for distinct channel subtypes in the presynaptic terminal do not exist (Lubbert *et al*., 2019b).

All in all, by demonstrating that motifs in the Ca_V_2 α_1_ subunit that control presynaptic Ca_V_2 subtype abundance are distinct from those determining presynaptic Ca_V_2 subtype preference, our findings provide new insights into the regulation of presynaptic Ca_V_2 subtype levels.

## Additional information section

### Author contributions

J.L. and P.V., carried out experiments and analyzed data. S.M.Y, Jr developed the mouse model S.M.Y, Jr planned the project and analyzed data. J.L. and S.M.Y, Jr. wrote the manuscript. All authors jointly revised the paper. All authors approved the final version of the manuscript, agree to be accountable for all aspects of the work in ensuring that questions related to the accuracy or integrity of any part of the work are appropriately investigated and resolved, all persons designated as authors qualify for authorship, and all those who qualify for authorship are listed.

## Acknowledgements

We thank members of the Young lab for their comments and critiques of the manuscript and project. We thank Hao Li for help with HdAd viral vector production. We thank Dr. Susan Stamnes and University of Iowa viral vector core, for help with HdAd vector production and tittering. We thank Dr. Phillip Ng for HdAd packing plasmids. We thank Dr. Chris Ahern and Miranda Schene for sharing HEK293T cells and HEK293T whole cell recording protocols. We thank Dr. Henry Colecraft for gifts of CMV Ca_V_β4 plasmid.

